# The metastable XBP1u transmembrane domain defines determinants for intramembrane proteolysis by signal peptide peptidase

**DOI:** 10.1101/322107

**Authors:** Sara Suna Yücel, Walter Stelzer, Alessandra Lorenzoni, Manfred Wozny, Dieter Langosch, Marius K. Lemberg

**Affiliations:** Centre for Molecular Biology of Heidelberg University (ZMBH), DKFZ-ZMBH Alliance, Im Neuenheimer Feld 282, 69120 Heidelberg, Germany; Center for Integrated Protein Science Munich (CIPSM) at the Lehrstuhl Chemie der Biopolymere, Technical University of Munich, Weihenstephaner Berg 3, 85354 Freising, Germany; MassMap GmbH & Co. KG, Meichelbeckstr. 13a, 85356 Freising, Germany

**Keywords:** Regulated intramembrane proteolysis, GxGD aspartic intramembrane protease, ERAD, exosite, subsite, transmembrane helix dynamics, XBP1u, HO1, SPP

## Abstract

Unspliced *XBP1* mRNA encodes XBP1u, the transcriptionally inert variant of the unfolded protein response (UPR) transcription factor XBP1s. XBP1u targets its mRNA-ribosome-nascent-chain-complex to the endoplasmic reticulum (ER) to facilitate UPR activation and prevents overactivation. Yet, its membrane association is controversial. Here, we use cell-free translocation and cellular assays to define a moderately hydrophobic stretch in XBP1u that is sufficient to mediate insertion into the ER membrane. Mutagenesis of this transmembrane (TM) region reveals residues that facilitate XBP1u turnover by an ER-associated degradation route that is dependent on signal peptide peptidase (SPP). Furthermore, the impact of these mutations on TM helix dynamics was assessed by residue-specific amide exchange kinetics, evaluated by a semi-automated algorithm. Based on our results, we suggest that SPP-catalyzed intramembrane proteolysis of TM helices is not only determined by their conformational flexibility, but also by side chain interactions near the scissile peptide bond with the enzyme’s active site.

## Introduction

The endoplasmic reticulum (ER) controls the synthesis, folding and modification of almost one third of our proteome. Under changing physiological circumstances or proteotoxic stress, the folding capacity of the ER may face an imbalance that can lead to an accumulation of misfolded proteins. In order to keep the ER functional and restore homeostasis, ER stress activates an intracellular signaling pathway called the unfolded protein response (UPR), which transcriptionally upregulates folding chaperones and the ER-associated degradation (ERAD) machinery (Walter and Ron, 2011). Subsequently, ER folding capacity increases and terminally misfolded protein species are targeted for proteasomal degradation by a diverse set of ERAD factors (Christianson and Ye, 2014). Chronic ER stress leads to UPR induced apoptosis, which emerges as a driver of several human diseases including diabetes and neurological disorders (Hetz and Papa, 2018).

In metazoans, three UPR signaling branches act in parallel; regulated intramembrane proteolysis of ATF6, translational control by PERK kinase and an IRE1-mediated unconventional splicing of the *XBP1* mRNA (Walter and Ron, 2011). IRE1 is a membrane integral stress sensor that oligomerizes upon accumulation of unfolded proteins in the ER lumen and cleaves XBP1 mRNA (Li et al., 2010; Yoshida et al., 2001). Consequently, the *XBP1* mRNA is spliced (removing a 26-nt intron) leading to an open reading frame that encodes for the mature transcription factor XBP1s, comprising an N-terminal basic leucine zipper (bZIP) dimerization domain followed by a C-terminal transcription activation domain. In contrast, the unspliced variant of *XBP1* mRNA encodes XBP1u, which consists of a hydrophobic stretch in the C-terminal portion and lacks the transcription activation domain. Although short lived, XBP1u acts as a negative regulator of the UPR by targeting XBP1s and activated ATF6 for degradation and thereby fine-tuning the UPR (Tirosh et al., 2006; Yoshida et al., 2006; Yoshida et al., 2009). Moreover, XBP1u mediates crosstalk between ER stress and the transcriptional control of autophagy (Zhao et al., 2013). Interestingly, newly synthesized XBP1u targets its own mRNA-ribosome-nascent chain complex to the ER in order to facilitate splicing of its mRNA by IRE1 (Yanagitani et al., 2009; Yanagitani et al., 2011). When emerging from the translating ribosome, the C-terminal hydrophobic stretch of XBP1u is recognized by the signal recognition particle (SRP) resulting in co-translational targeting to the Sec61 translocon (Kanda et al., 2016; Plumb et al., 2015). Different to insertion of canonical nascent signal peptides and signal anchor sequences (Mothes et al., 1994; Voorhees and Hegde, 2016), Kohno and co-workers suggest that XBP1u interacts with Sec61 and the ER membrane on the cytosolic side, from where it may detach to reach the nucleus in an unprocessed form (Kanda et al., 2016; Yanagitani et al., 2009). In contrast, our previous research indicated that the XBP1u hydrophobic region is capable of inserting into the ER membrane as a type II transmembrane (TM) domain, having its C-terminus in the ER lumen (Chen et al., 2014). Furthermore, we showed that a truncated version of XBP1u, where the 54 amino acid C-terminal tail has been deleted, is inserted into the ER membrane as a tail-anchored (TA) protein (Chen et al., 2014). This indicates that the GET pathway, responsible for post-translational targeting and insertion of TA proteins (Hegde and Keenan, 2011), and the co-translational targeting machinery, both are capable of integrating XBP1u’s hydrophobic domain into the ER membrane. However, XBP1u is subsequently cleaved by signal peptide peptidase (SPP) and targeted for proteasomal degradation (Chen et al., 2014). SPP belongs to a special group of enzymes, so-called intramembrane proteases that cleave TM anchors of a wide range of proteins to trigger their release from cellular membranes (Lemberg, 2011). Intramembrane proteases comprise aspartic proteases such as SPP and γ-secretase; rhomboid serine proteases; S2P metalloproteases and the glutamyl protease Rce1. They are mechanistically different, but all form membrane-embedded aqueous active sites and face similar problems in substrate recognition, unwinding of substrate TM helices and cleavage (Langosch et al., 2015; Strisovsky, 2016; Wolfe, 2009).

While SPP was initially identified as the protease clearing signal peptides from the ER membrane (Lemberg and Martoglio, 2002; Weihofen et al., 2002), it is now known that SPP also cleaves certain *bona fide* TM domains (Boname et al., 2014; Chen et al., 2014; Hsu et al., 2018; Hsu et al., 2015). This indicates that there are mechanistic parallels to SPP-like (SPPL) proteases, which cleave type II membrane proteins in the Golgi and the late secretory pathway (reviewed by (Mentrup et al., 2017)). However, to target ERAD substrates for degradation, SPP interacts with the rhomboid pseudoprotease Derlin1 and the E3 ubiquitin ligase TRC8, forming a 500-kDa complex (Chen et al., 2014; Stagg et al., 2009). While TRC8 ubiquitinates XBP1u to ensure its efficient turnover by the proteasome (Boname et al., 2014; Chen et al., 2014), Derlin1 acts as a substrate receptor for the luminal domain of XBP1u (Chen et al., 2014). This function of Derlin1 obviates the need for signal peptidase cleavage of this substrate, as required for SPP-catalyzed cleavage of signal peptides (Lemberg and Martoglio, 2002). In absence of Derlin1, XBP1u is only recognized by SPP as a TA variant (Chen et al., 2014). Although the list of substrates and the physiological significance of SPP in the cell is expanding, its substrate recognition mechanism and cleavage determinants are still ill-defined.

Efficient SPP-catalyzed intramembrane cleavage of signal peptides requires amino acid residues predicted to lower TM helix stability (Lemberg and Martoglio, 2002). In line with this, our previous analysis of XBP1u showed that its SPP-mediated cleavage depends on polar side chain residues as mutating those into leucine prevent proteolysis and lowered the helicity profile in CD spectroscopy (Chen et al., 2014). However, it remains unclear whether SPP recognizes a certain consensus sequence as has been described for bacterial rhomboid intramembrane serine proteases (Strisovsky et al., 2009). Here, we elucidate the recognition of XBP1u by the Sec61 translocon and show that the moderately hydrophobic segment of XBP1u is sufficient for membrane insertion. We use a combination of *in vitro* translocation and cleavage assays, cell-based assays and deuterium/hydrogen exchange (DHX) experiments to understand the effects of conformational dynamics in the TM helix and the influence on substrate recognition and cleavage by SPP. Most significantly, we find that SPP-catalyzed cleavage of *bona fide* TM helices not only requires conformational flexibility in the N-terminal TM segment but also specific side chain interactions.

## Results

In order to characterize the efficiency of membrane insertion of XBP1u and to define requirements for its cleavage by SPP, we i) examined the role of its hydrophobic region in cell-based and *in vitro* assays and ii) correlated the impact of various mutations within this TM domain on helix dynamics with efficiency of SPP-mediated cleavage and degradation.

### Turnover of XBP1u is governed by its TM domain

Our previous investigation revealed that XBP1u, subsequent to ER insertion as a type II membrane protein, is turned over by an SPP-dependent ERAD pathway (Chen et al., 2014). In the context of ERAD, XBP1u is recognized in the luminal portion by Derlin1 priming it for SPP-mediated turnover circumventing trimming by signal peptidase or a short juxtamembrane C-terminus of a TA protein (Chen et al., 2014). However, the TM determinants for SPP-catalyzed XBP1u cleavage had not been investigated in detail. One way to interfere with SPP-mediated degradation is to exchange hydrophilic TM residues by leucine (Lemberg and Martoglio, 2002). Earlier, we have shown that mutating the N-or the C-terminal TM portion referred to mt1 and mt2, respectively (Figure 1A), blocks SPP-catalyzed cleavage of XBP1u (Chen et al., 2014). To test the ER insertion orientation and efficiency of these TM domain variants we took advantage of an established XBP1u-R232N glycosylation reporter construct in tissue culture cells (Chen et al., 2014). Both mutants promoted efficient insertion as type II membrane proteins, even with a higher efficiency than wt (Figure 1B). Increasing the hydrophobicity (Table S1) of the XBP1u TM region results in a more stably integrated fraction into the ER in a type II membrane topology orientation.

**Figure 1.**
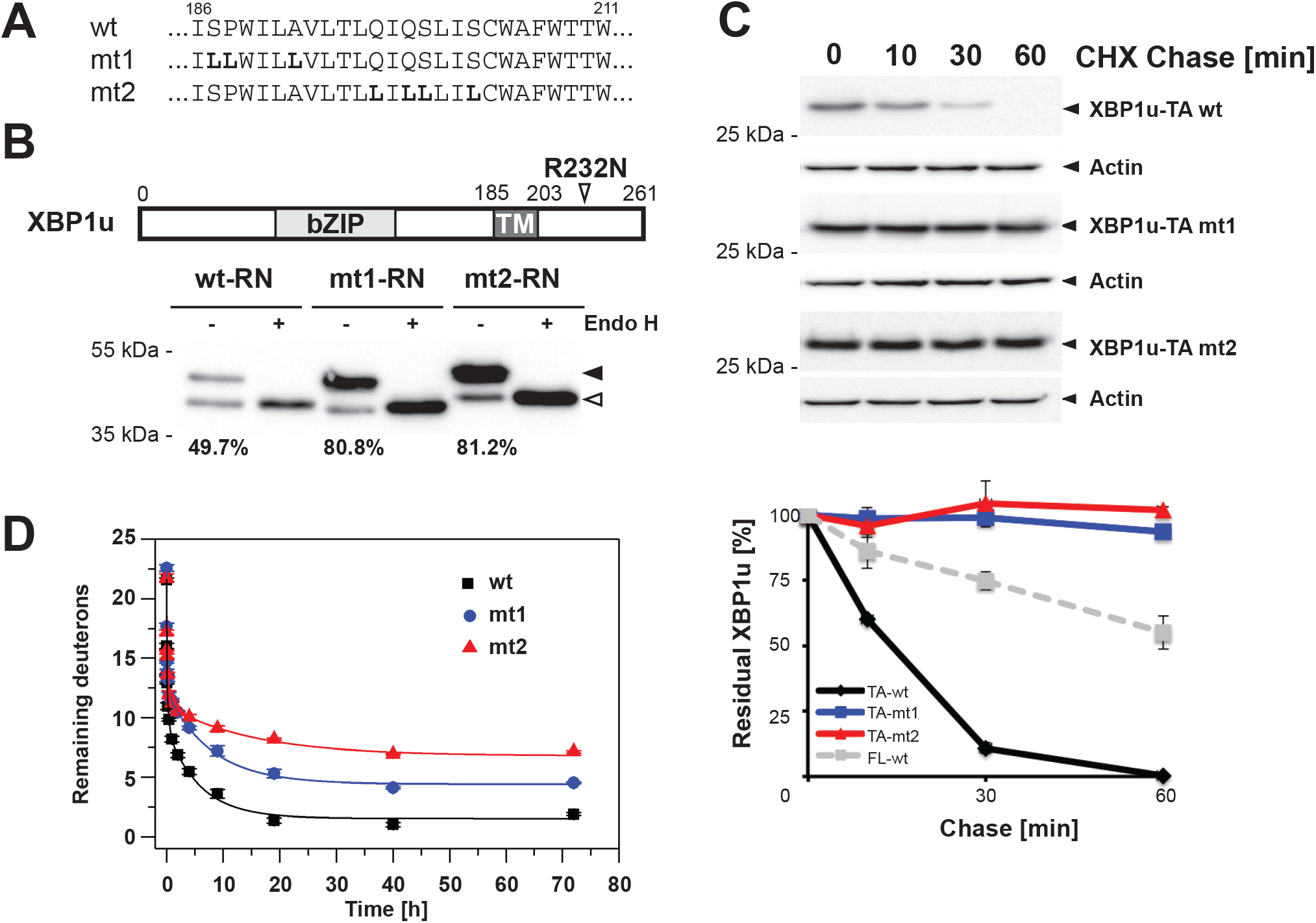
Turnover of XBP1u is governed by its TM domain. **(A)** Sequences of XBP1u TM domain leucine mutants on the N-terminal (mt1) and on the C-terminal (mt2). TM domains are underlined. **(B)** XBP1u wt, mt1 and mt2 ER targeting and insertion as type II membrane protein assessed by Endo H-sensitivity of the glycoslylation mutant R232N (RN) (triangle). Hek293T cells are transfected with FLAG-tagged XBP1u-RN harboring wt, mt1 or mt2 TM domain and cell lysates are treated with EndoH. Filled arrowhead, glycosylated protein; open arrowhead, unglycosylated protein. Glycosylation efficiencies are indicated respectively and were calculated as the intensity of the glycosylated band divided by the total. **(C)** FLAG-XBP1u-TA mt1 and mt2 are stable, whereas the wt construct is rapidly degraded as assessed by cycloheximide (CHX) chase and western blotting using antiFLAG antibody (mean ± SEM, n=3). Quantification of the degradation kinetics of the full-length (FL) XBP1u is shown for comparison. **(D)** Overall amide DHX kinetics of XBP1u wt TM domain compared to mt1 and mt2 (n ≥ 3, SEM are smaller than the sizes of the symbols). All used peptides contain N-and C-terminal KKK tags for better solubility and gas phase fragmentation.

In order to uncouple SPP-mediated cleavage from Derlin1-dependent recruitment, we created TA-variants of XBP1u, which lack the C-terminal 54 amino acids. We assessed the degradation kinetics of XBP1u-TA wt, mt1 and mt2 in a cycloheximide chase experiment in transfected tissue culture cells. The half-life of XBP1u-TA wt diminished to approximately 15 minutes compared to 60 minutes of the full-length XBP1u construct consistet with our previous report (Figure 1C) (Chen et al., 2014). In contrast, XBP1u-TA mt1 and mt2 were both stable and no decay was observed over the chase time (Figure 1C), while ER localization was not affected as assessed by immunofluorescence microscopy (Figure S1A). This indicates that the TM region of XBP1u in the context of a tail-anchored protein determines the rate of SPP-dependent proteolysis.

### Increasing rigidity of TM helix prevents SPP-catalyzed cleavage

Next, we investigated the effects of these XBP1u mutations on the helicity and the conformational flexibility of the TM domain by amide DHX kinetics. To this end, we created synthetic peptides and dissolved them in aqueous trifluoroethanol (TFE), which serves as a mimic of the aqueous environment within the catalytic cleft of an intramembrane protease and readily solubilizes the peptides (Pester et al., 2013). CD spectroscopy confirmed that the peptides form >60% helix in this solvent, with a small but significant increase in the helicity of mt1 (Figure S1B), as shown before (Chen et al., 2014). This ensures that the following measurements predominantly describe the properties of the helical state. The overall conformational flexibility of exhaustively (> 95%) deuterated peptides was assessed in the same solvent by recording DHX kinetics that reflects the stability of intrahelical amide hydrogen-bonds (H-bonds). At time point zero, where the labile deuterons bound to non-H-bonded polar atoms have already exchanged for protons, ~22 deuterons remain (Figure 1D). These remaining deuterons represent >90% of those 24 backbone amides that can at least be partially protected from DHX by intrahelical H-bond formation. As exemplified by the wt spectra shown in Figure S1C, a gradual shift in addition to the lack of a bimodal shape of the isotopic distributions is diagnostic of transient local unfolding events described by a kinetic model, known as EX2 kinetics, where individual deuterons exchange in an uncorrelated fashion (Konermann et al., 2011). Next, we investigated the leucine substitutions in the cleavage-deficient mt1 and mt2. Leucine substitutions are predicted to stabilize intrahelical contacts between their large and flexible side chains and the side chains of other residues within the helix (Quint et al., 2010). Consistent with this, amide exchange of mt1 and mt2 was slower than that of wt, indicating their lower average helix backbone flexibility (Figure 1D). This refines our previous structural analysis solely relying on CD spectroscopy (Figure S1B and (Chen et al., 2014)) in revealing that mutagenizing either N-or C-terminal TM residues to leucine render the TM helix more rigid. Together, these results corroborate that the XBP1u TM segment serves as the key determinant for SPP-catalyzed intramembrane proteolysis.

### Interaction of XBP1u with the Sec61 translocon leads to insertion of a TM segment

ER targeting of XBP1u has been described to be in an SRP-dependent pathway (Kanda et al., 2016). However, to which extend it inserts into the Sec61 translocon and whether it adopts a type II membrane anchor is still controversially debated (Chen et al., 2014; Kanda et al., 2016; Plumb et al., 2015). Our previous analysis of the full-length XBP1u protein in cells and an *in vitro* translocation assay suggested that a large fraction adopts a type II topology (Figure 1B and (Chen et al., 2014)). Here, we choose to analyze the XBP1u TM region in isolation to decipher its influence on the functional interaction with the Sec61 translocon.

First, we analyzed the hydrophobic region of XBP1u based on a thermodynamic scale that predicts membrane insertion efficiency of TM segments, which had been experimentally determined by von Heijne and colleagues (Hessa et al., 2007). According to this scale, the apparent free energy (ΔG_app_) of XBP1u is not predicted to form a stable TM segment compared to the thermodynamically favorable insertion of a *bona fide* TM helix such as calnexin (ΔG_app_ < 0 kcal/mol) (Figure 2A). However, single-spanning membrane proteins containg a moderately hydrophobic TM domain segment (ΔG_app_ > 0 kcal/mol) are prevalent and lead to co-translational membrane insertion (De Marothy and Elofsson, 2015; Dou et al., 2014; Junne and Spiess, 2017). According to this prediction algorithm signal sequences also frequently have positive ΔG_app_ values (see Figure 2A for examples) and were therefore excluded in the analysis by von Heijne (Hessa et al., 2007). Nonetheless, it is well accepted that the Sec61 translocon mediates efficient insertion of signal peptides of nascent peptide chains (Mothes et al., 1994; Voorhees and Hegde, 2016), indicating that the ΔG_app_ prediction tool does not reflect the full scale of TM segments.

**Figure 2.**
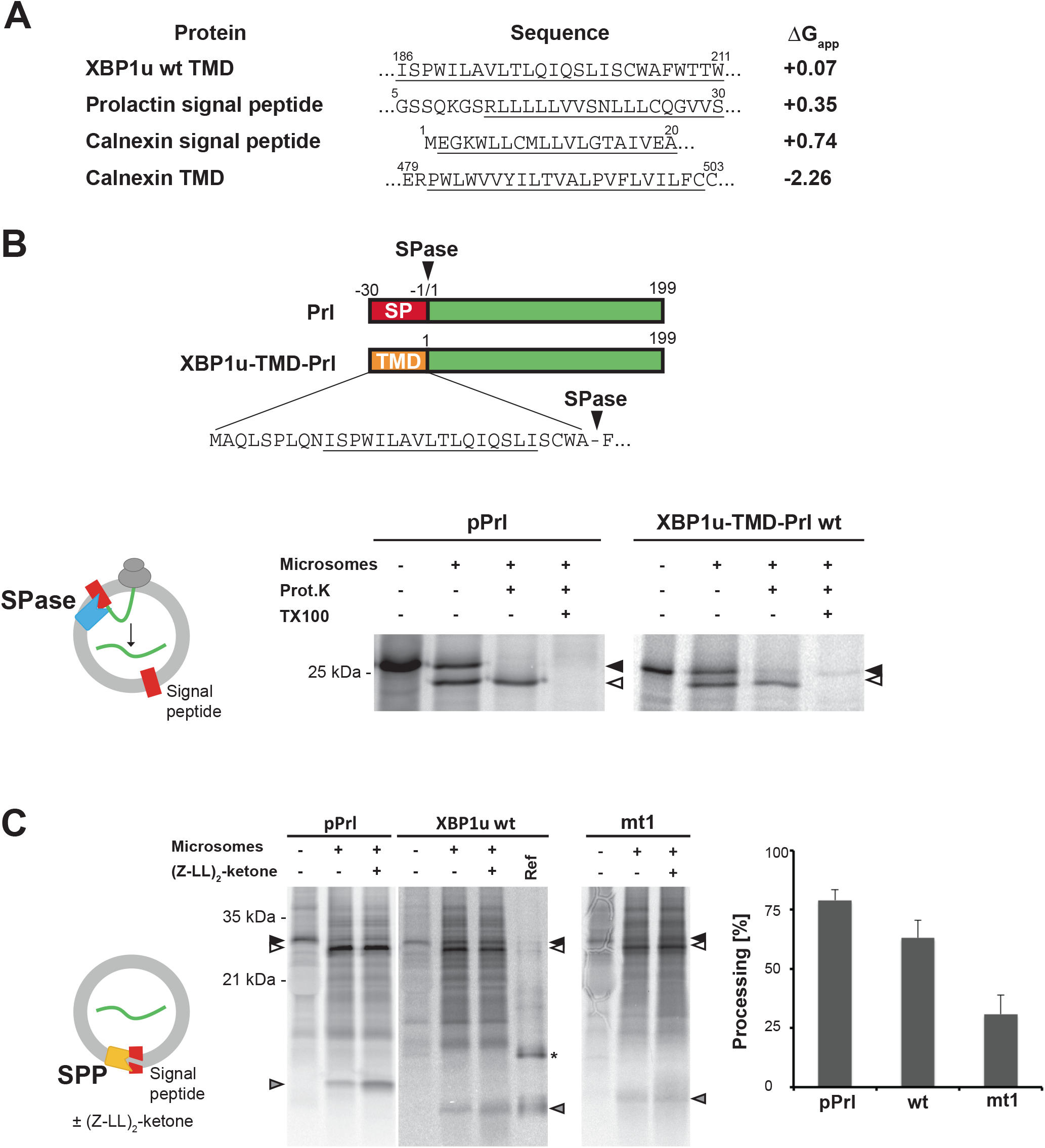
XBP1u TM domain is sufficient for insertion into the ER membrane in a type II orientation. **(A)** Predicted apparent free energy differences (ΔG_app_) for Sec61 insertion of the XBP1u TM domain (TMD) and ER-targeting signal peptides (SP) compared to the TMD of calnexin (Hessa et al., 2007). Sequences taken for the calculation are underlined. **(B)** Translocation and protease protection assay revealing that the XBP1u TM domain is sufficient to mediate ER insertion. Upper: Schematic representation of Prl and XBP1u-TMD-Prl constructs. Lower left: Illustration of the cell-free *in vitro* translation/translocation assay. Upon the addition of microsomes, the pre-protein is translocated and the signal sequence (red) is cleaved of by signal peptidase (blue) to release the mature protein (green). Lower right: *In vitro* translation of mRNA encoding either Prl or the XBP1u-TMD-Prl fusion proteins in the absence or presence of canine rough microsomes. Reactions were treated with Proteinase K and Triton X-100 as indicated and analyzed by SDS-PAGE and autoradiography. Filled arrowhead, pre-protein; open arrowhead, translocated Prl. **(C)** Processing of signal peptides in microsomes. Left: Illustration of microsome-based *in vitro* SPP assay (represented in yellow) leading to cleavage and release of membrane-spanning signal peptide (red). Middle: *In vitro* translation of mRNA encoding pPrl, XBP1u-TMD-Prl fusion proteins (wt and mt1) in the absence or presence of ER-derived microsomes and 10 μM of (Z-LL)2-ketone. Right: Quantification of signal peptide processing comparing relative amounts of signal peptide obtained from vehicle only to (Z-LL)2-ketone treated sample (mean ± SEM, n=4). Ref, shows a reference peptide comprising the XBP1u-TMD neo-signal peptide; asterisk, putative dimer of *in vitro* translated reference signal peptide.

We therefore asked whether the putative TM region of XBP1u interacts with the Sec61 translocon in the manner of a nascent signal peptide. Signal peptides have a common three-domain structure comprised by a short positive-charged ‘n-region’, a hydrophobic ‘h-region’ of 7 to 15 residues and a polar ‘c-region’ including the site for signal peptidase cleavage (Haeuptle et al., 1989). While n-regions are commonly only few amino acids long, internal signal sequences with protrusions in the size range of functional protein domains have also been reported (Kapp et al., 2009). With a low scoring signal peptidase cleavage site predicted by SignalP 3.0 (Bendtsen et al., 2004) in the putative c-region between alanine-206 and phenylalanine-207 (Figure 2B), the XBP1u hydrophobic region shows similarity to an internal signal sequence. Hence, we created a chimeric protein in which we replaced the signal sequence of pre-prolactin (pPrl) with this predicted XBP1u TM domain and N-terminal flanking residues (XBP1u-TMD-Prl) and tested whether this XBP1u “neo-signal peptide” is sufficient to mediate ER targeting and translocation in a microsome-based cell-free system (Figure 2B). To this end, mRNA encoding either pPrl wild-type (wt) or the XBP1u-TMD-Prl fusion was translated in reticulocyte lysate supplemented with [^35^S]-methionine and cysteine. Upon addition of ER-derived canine rough microsomes, signal sequences are inserted into the membrane leading to signal peptidase cleavage and liberation of the translocated polypeptide (Blobel and Dobberstein, 1975). Indeed, a faster migrating band co-migrating with mature Prl was observed for XBP1u-TMD-Prl, indicating that in the context of the XBP1u-TMD-Prl fusion protein the cryptic signal peptidase site is processed. This product was protected from proteinase K but became proteinase K-sensitive when detergent was added to solubilize the microsomal membrane (Figure 2B). These results show that the XBP1u TM domain alone is sufficient for productive interaction with Sec61 and subsequent translocation of a downstream fusion protein. Furthermore, the N-terminal variant (mt1) promotes efficient translocation of Prl. Despite a predicted negative ΔG_app_ of the mt2 TM domain, the neo-signal sequence of the Prl fused version (XBP1u-TMD-Prl mt2) did not promote efficient ER insertion as a type-II oriented signal sequence (Figure S2 and Table S1). Low amount of a protease protected form may indicate that the construct leads to a type II signal anchor, yet at low efficiency.

To test *in vitro* for SPP-catalyzed processing of the neo-signal peptide, i.e. the XBP1u TM segment, we isolated microsomes from a co-translational translocation assay using pPrl, XBP1u-TMD-Prl wt and mt1 mRNA. Upon cleavage of the pre-protein by signal peptidase, liberated signal peptides are anchored in the microsomal membrane similar to a membrane protein with a single TM region until they are released by SPP-catalyzed intramembrane proteolysis (Lemberg et al., 2001; Weihofen et al., 2000). We quantified SPP activity by isolating microsomes from the reaction in the presence of the SPP inhibitor (Z-LL)2-ketone and compared the relative amount of signal peptides obtained in inhibitor treated membranes with the vehicle control (Figure 2C) (Weihofen et al., 2000). As had been observed for pPrl, also for the liberated XBP1u neo-signal peptide, a weak band in the low molecular range emerged, which is stabilized upon treatment with (Z-LL)2-ketone (Figure 2C). This product co-migrated with an *in vitro* translated reference peptide comprising the XBP1u neo-signal peptide. The corresponding signal peptide derived from XBP1u-TMD mt1, however, was only processed by approximately 30% compared to >60% for the XBP1u wt TM domain sequence (Figure 2C). This reduction of SPP processing efficiency is in the range observed for signal peptides lacking obvious helix-destabilizing residues (Lemberg and Martoglio, 2002).

### Increasing TM helix flexibility has differential effects on cleavage

Results described above provide further evidence that XBP1u spans the ER membrane with a metastable type II TM anchor that is rapidly processed by SPP to target it for proteasomal degradation (Chen et al., 2014). We next asked how increased flexibility of the TM helix modulates its cleavage by SPP. To this end, we inserted a di-glycine motif in the N-terminal (mt3), central (mt4) or C-terminal (mt5) portion of the TM domain (Figure 3A). Due to the absence of a side chain, glycine destabilizes a TM helix by introducing a packing defect (Hogel et al., 2018).

**Figure 3.**
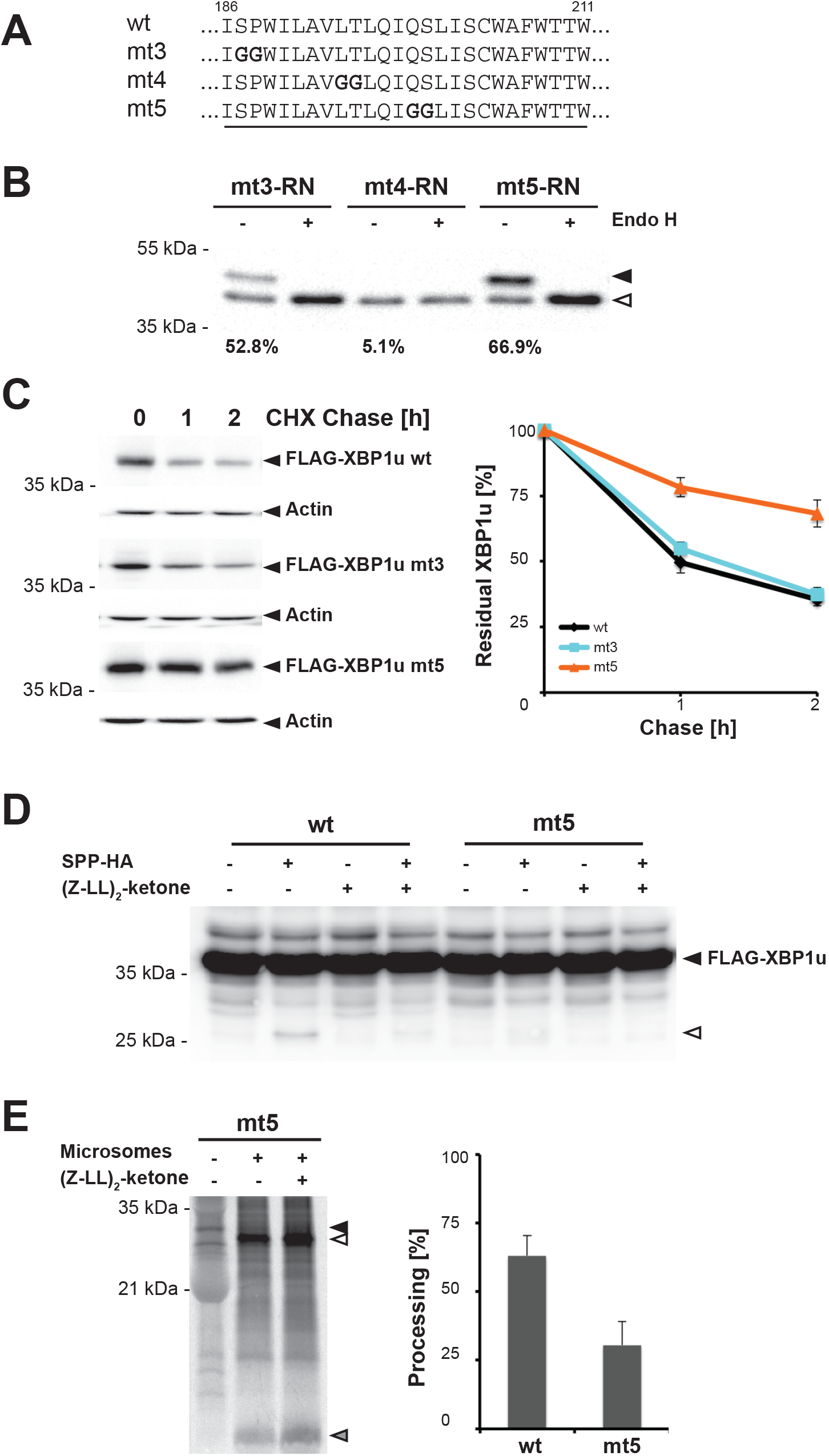
Positional effect of di-glycine motifs on SPP-catalyzed turnover. **(A)** Sequences of XBP1u di-glycine TM domain mutants. **(B)** ER targeting and insertion efficiencies of XBP1u mt3, mt4 and mt5 assessed by the R232N glycosylation reporter (RN). Mt3 and mt5 insert into the ER in a type II orientation whereas mt4 predominantly stays cytoplasmic. Filled arrowhead, glycosylated protein; open arrowhead, unglycosylated protein. Glycosylation efficiencies are indicated respectively. **(C)** Degradation kinetics of FLAG-XBP1u wt, mt3 and mt5 in transfected Hek293T cells as assessed cycloheximide (CHX) chase and western blotting using anti-FLAG antibody (mean ± SEM, n=3). **(D)** XBP1u cleavage assay in the presence of the proteasome inhibitor epoxomicin (1 μM). Hek293T cells were co-transfected with either FLAG-XBP1u wt or mt5, with empty vector or SPP-HA. Cell were treated either with vehicle or SPP inhibitor, 50 μM (Z-LL)2-ketone. Filled arrowhead, full-length XBP1u; open arrowhead, cleavage fragment created by SPP-mediated cleavage. **(E)** *In vitro* signal peptide cleavage of XBP1u-TMD-Prl mt5. Quantification of signal peptide processing comparing relative amounts of signal peptide obtained from vehicle only to (Z-LL)2-ketone treated sample (mean ± SEM, n=4). Quantification of XBP1u-TMD-Prl wt is shown for comparison (see Figure 2D).

First, we assessed the topology of the di-glycine mutants via the XBP1u-R232N glycosylation reporter. We observed that mt3 and mt5 are efficiently targeted to the ER and adopt a type II topology, whereas only a negligible fraction was glycosylated in the case of mt4 (Figure 3B). Consistent with this, immunofluorescence analysis revealed for mt3 and mt5 ER localization in the same range as observed for XBP1u wt, mt1 and mt2 (Figure S3A). In contrast, only a minor pool of mt4 co-localized with the ER marker and the predominant fraction was detected in the nucleus. Therefore, introducing the di-glycine motif into the center of the TM helix has the strongest effect on membrane insertion, as also reflected by a positive ΔG_app_ (Table S1), thus limiting the biological relevance of mt4. Next, we determined the stability of these mutants in transfected Hek293T cells by adding cycloheximide to block synthesis of new proteins and chase the pre-existing fraction of XBP1u over time. Interestingly, mt5 was resistant to degradation whereas mt3 showed degradation rates similar to XBP1u wt (Figure 3C). Likewise, the ER targeting-deficient mt4 showed not significant stabilization (Figure S3B), indicating that cytoplasmic/nucleoplasmic degradation pathways can also efficiently break down XBP1u. On the other hand, mt3, which is ER resident (Figure S3A) and adopts a type II topology (Figure 3B), is turned over at the same rate as the wt construct (Figure 3B). The degradation of mt3 is sensitive to (Z-LL)2-ketone (Figure S3C), indicating that, like XBP1u wt, it is a substrate for SPP-dependent ERAD. In contrast, mt5 accumulates in the ER and is stable over the entire chase time (Figure S3A and C). To detect potential XBP1u cleavage fragments we co-expressed XBP1u and SPP in the presence of the proteasome inhibitor epoxomicin. Although inhibition of the proteasome decreases also SPP activity and only low amounts of XBP1u cleavage fragment can be detected (Chen et al., 2014), upon ectopic expression of SPP we could observe an SPP-generated N-terminal cleavage fragment of XBP1u wt that is sensitive to (Z-LL)2-ketone (Figure 3D). In contrast, for mt5 only traces of this SPP-generated cleavage fragment was observed (Figure 3D). This strengthens the results from the cycloheximide chase assay showing that the protein is stabilized over time. Similarly, we observed inefficient SPP processing of the mt5 neo-signal peptide in the *in vitro* translocation assay (Figure 3E), corroborating that the C-terminal di-glycine motif hampers cleavage by SPP.

In order to detect a potential correlation between cleavability and helix flexibility, we determined amide exchange kinetics. As predicted, mt3, mt4 and mt5 exhibit faster overall DHX kinetics compared to the wt TM domain (Figure 4A). CD spectroscopy, however, showed similar overall helicity of the wt and mutants (Figure S4A), demonstrating that DHX is a much more sensitive tool to detect changes in helix flexibility (Figure 4A). A close inspection of the kinetics reveals the following rank order of the glycine impact: mt4 > mt5 > mt3 (Figure 4A, inset); this reflects the relative importance of the mutated positions for overall helix flexibility. To refine our analysis, we sought to correlate cleavability of our constructs with TM flexibility, which was resolved at the single-residue level. To this end, we performed electron transfer dissociation (ETD) of our peptides in the gas phase after different periods of DHX. Using a novel software solution that accelerates ETD fragment analyses by an order of magnitude, we determined the DHX kinetics of individual backbone amides that is quantitated by the exchange rate constants kexp shown in Figure S4B and Table S2. Since only incomplete kinetics could be determined for several N- and C-terminal residues due to rapid exchange near the helix termini, we focused on DHX kinetics of amides within the core of the XBP1u TM helix from isoleucine-186 to phenylalanine-207 (Figure S4C). At all these residues, the kexp values are below the respective calculated chemical amide exchange rate constants kch, which reflects their participation in secondary structure, *viz.* helix. Exchange rate constants were then converted to the free energy changes associated with H-bond formation ΔG (Table S2). Figure 4B shows the distribution of ΔG values for the wt TM helix that increase from ~2 kcal/mol near the helix N-terminus to ~4.5 kcal/mol. These values are plausible, given that values of ~1 kcal/mol were previously reported for intrahelical H-bonds in water, which can effectively compete with intrahelical H-bonding, ~4 kcal/mol for H-bonds of model compounds in apolar solvent, and 6.6 kcal/mol for H-bonds in vacuo (Bowie, 2011). To quantitate the effects of the mutations, we calculated the differences in H-bond stability ΔΔG by subtracting wt ΔG values from the respective mutant values. The results (Figure 4C) reveal that mutations to leucine in mt1 and mt2 stabilize local amide H-bonds by up to ~3 kcal/mol; in case of mt1, stabilization extends to several residues downstream of the mutations. By contrast, the di-glycine mutations in mt3 and mt5 destabilize by up to ~1 kcal/mol (Figure 4C). For mt4, which fails efficient ER targeting, the extent of destabiliziation is much larger with up to ~4 kcal/mol around the central di-glycine motif (Figure S4D). Taken together, these results illustrate that introducing leucine or glycine residues into the XBP1u TM domain has profound effects on TM helix flexibility, but there is no clear correlation to cleavability by SPP. Contrary to the initial proposal that SPP cleavage efficiency is primarily determined by a rate-limiting unfolding of the substrate TM helix into the active site (Lemberg and Martoglio, 2004), mutations destabilizing the TM domain (mt5) can also reduce SPP processing efficiency.

**Figure 4.**
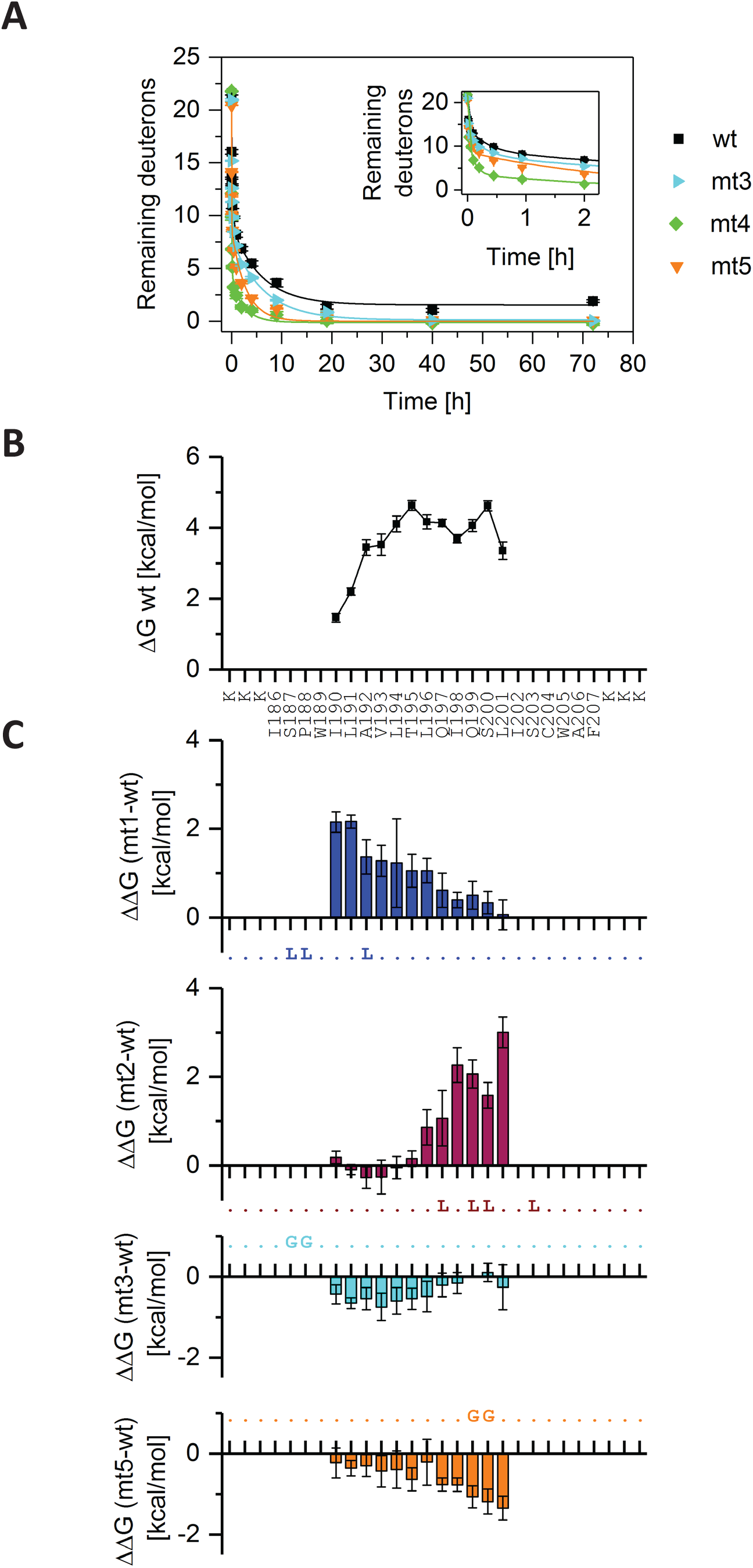
Di-glycine mutants show local TM helix destabilization. **(A)** Overall amide DHX kinetics of XBP1u wt compared to mt3, mt4 and mt5 recorded from the masses of triply charged ions (n > 3, SEM are smaller than the sizes of the symbols). The inset focuses on the initial 2 h of the incubation period. **(B)** Free energy change ΔG of intrahelical amide H-bond formation. **(C)** Differences in ΔG of wt and ΔG of mutant TM domains (ΔΔG). Values are only shown for those residues where sufficient data points could be collected for wt and the respective mutant (see Table S2). Wt residues in the mutant TM domain sequences are represented as dots. Error bars correspond to standard confidence intervals calculated from the standard errors of kexp.

### Single residues can affect cleavage

Since the C-terminal di-glycine motif in XBP1u mt5 prevents SPP-dependent degradation (Figure 3C and D), cleavage by SPP may not only depend on TM helix dynamics but also on substrate recognition requiring specific residues. In order to study the importance of glutamine-199 and serine-200 that were simultaneously mutated in mt5, we generated two single glycine mutants, namely Q199G and S200G (Figure 5A). We confirmed ER targeting and topology of these single TM domain mutants by the glycosylation reporter assay (Figure S5A) and immunofluorescence analysis (Figure S5B). We tested their degradation rate by cycloheximide chase experiments (Figure 5B and S5C). Interestingly, the S200G mutant was stabilized to similar levels as mt5, while Q199G showed a degradation rate comparable to XBP1u wt, suggesting that SPP recognizes certain side chains such as serine-200. This observation was supported in a semi-quantitative SPP cleavage assay, where a cleavage fragment for the XBP1u wt and the Q199G mutant was detected but to less extent for the S200G mutant (Figure 5C). Therefore, we wondered whether other residues in the C-terminal portion of the XBP1u TM domain also contribute to the specificity of SPP-mediated degradation. We performed a glycine scan in which we mutated each residue starting from leucine-196 to cysteine-204 into a glycine and tested their degradation rate by cycloheximide chase experiments (Figure 5A, D and S5D). Interestingly, the XBP1u glutamine-197 (Q197G) mutant as well was stabilized, while the other mutations did not stabilize XBP1u. This indicates that certain polar residues in the C-terminal portion of XBP1u play a distinct and important role for SPP-mediated recognition and cleavage.

**Figure 5.**
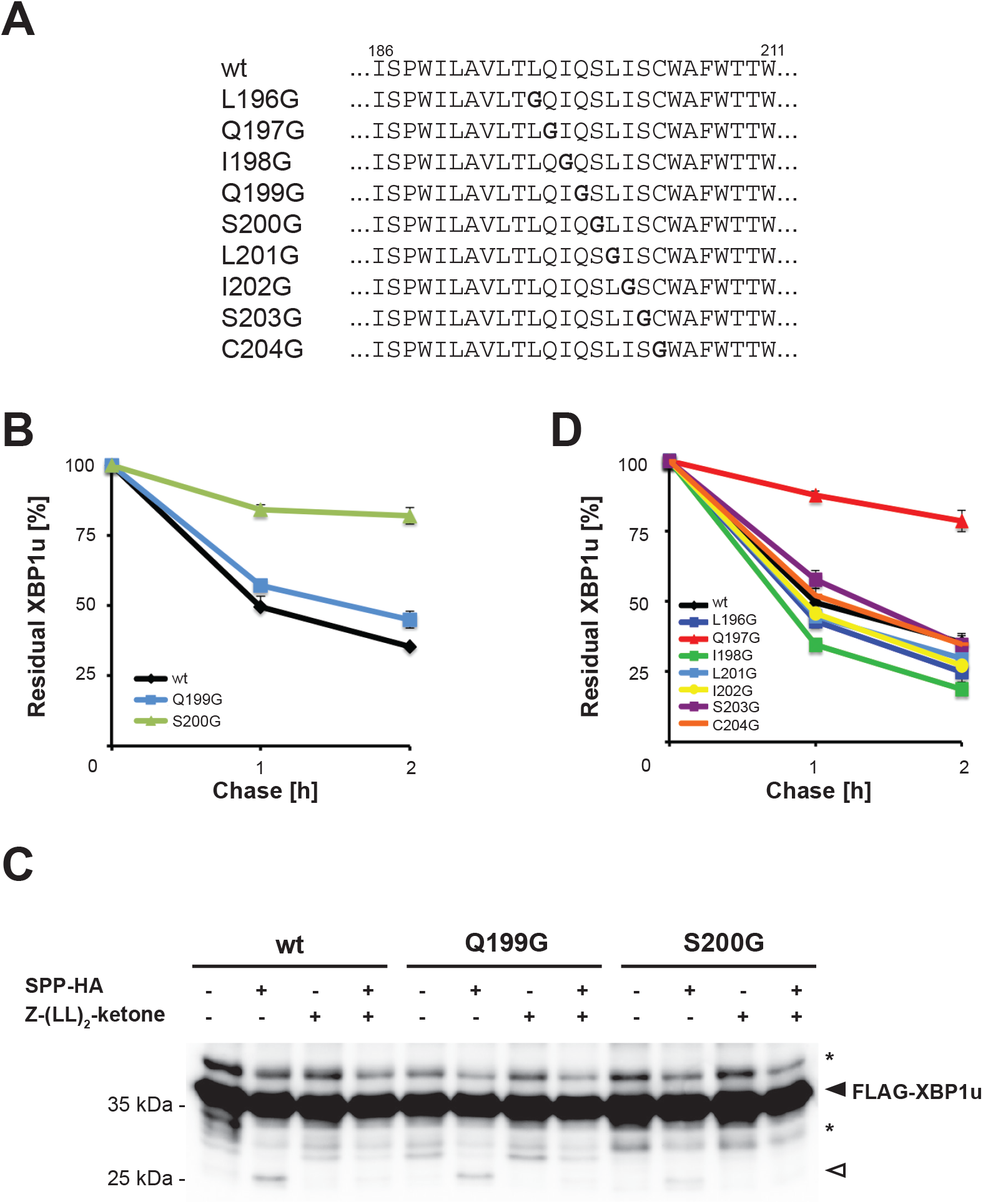
XBP1u turnover depends not only on global TM helix dynamics but also on residue specific features. **(A)** Sequences of XBP1u single-residue TM domain mutants. **(B)** Degradation kinetics of FLAG-XBP1u Q199G and S200G (see Figure S5C for western blots). Quantification of XBP1u wt is shown for comparison (see Figure 3C). **(C)** XBP1u cleavage assay in the presence of the proteasome inhibitor epoxomicin (1 μM). Hek293T cells were co-transfected with FLAG-XBP1u wt, Q199G or S200G mutants, either with empty vector or SPP-HA. Cell were treated either with vehicle or SPP inhibitor, 50 μM (Z-LL)_2_-ketone. Filled arrowhead, full-length XBP1u; open arrowhead, cleavage fragment created by SPP-mediated cleavage. **(D)** Glycine scan of the C-terminal portion of XBP1u TM domain. Degradation kinetics of FLAG-XBP1u mutants assessed by CHX chase and western blotting (see also Figure S5D).

We note, that several well-characterized SPP substrates such as the signal peptides of calreticulin or insulin do not have a serine or a glutamine in the respective position (Kronenberg-Versteeg et al., 2018; Lemberg and Martoglio, 2002). Despite that, we wondered whether this features are important to foster processing of actual TM helices. Based on a recent proteomics screen, the TA protein heme oxygenase 1 (HO1) has been shown to be a native substrate for the SPP-dependent ERAD pathway (Boname et al., 2014), but its substrate determinants have not been analyzed yet. Remarkably, HO1 has in addition to potential helix destabilizing features (proline-266 and arginine-269) in the N-terminal TM domain portion also a serine-275 in the C-terminal half (Figure 6A). To test the influence of these putative substrate determinants, we mutated them individually to leucine or alanine and tested their steady state levels (Figure 6B). Our rationale was to stabilize the N-terminal TM domain portion by leucine and to remove putative side-chain interactions in the C-terminal half by inserting alanine (without introducing further flexibility as may be observed for glycine). All HO1 mutants tested had an approximately three-fold steady state increase compared to wt (Figure 6B) and showed a reduced degradation kinetics in a cycloheximide chase (Figure 6C and S6A). These results indicate that like for XBP1u, dynamics of the N-terminal TM helix portion and a serine in the C-terminal half play a role in efficient recognition of HO1 by SPP.

**Figure 6.**
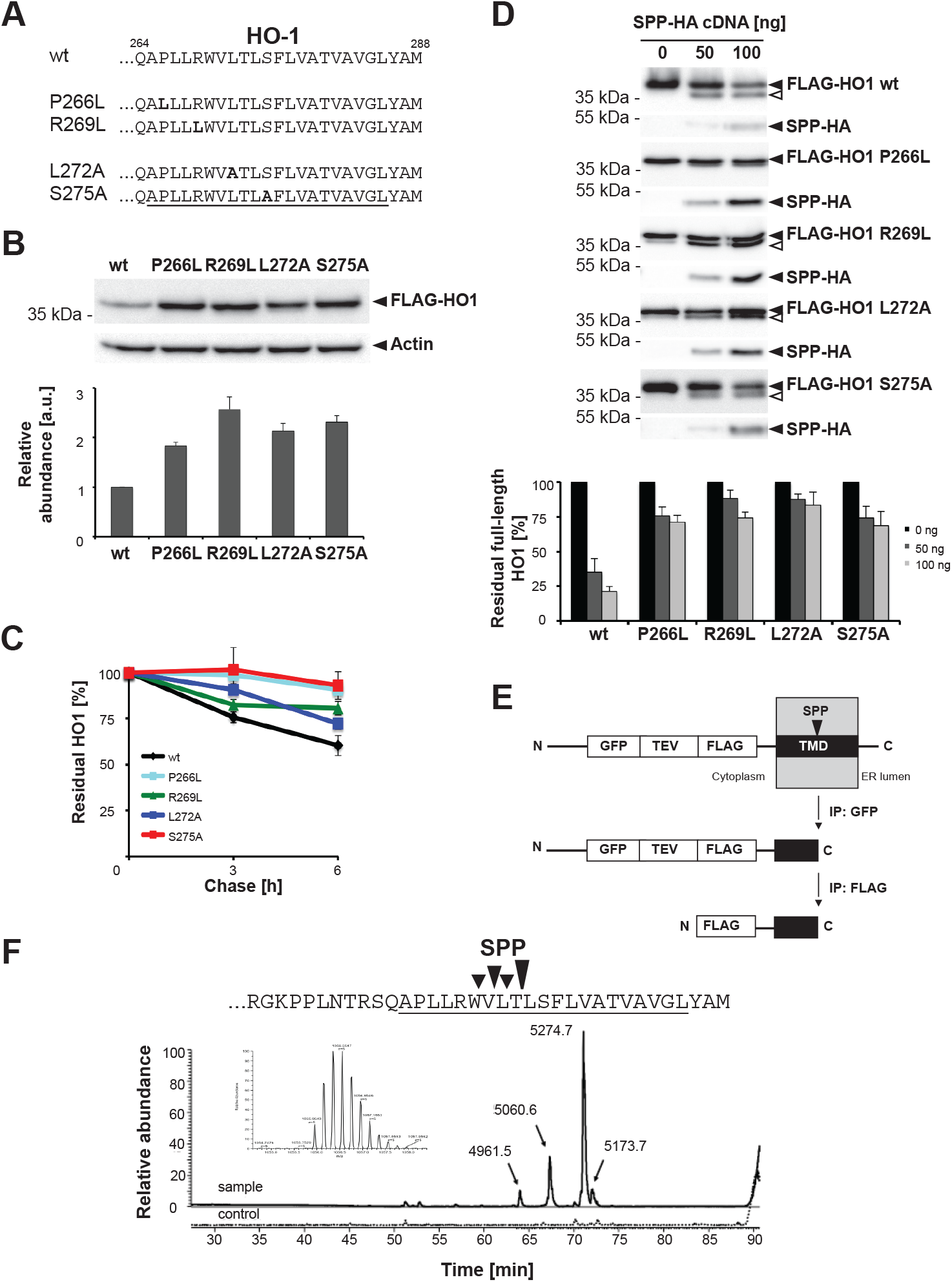
Cleavage site motif facilitates SPP-catalyzed cleavage of TA proteins. **(A)** Sequence of HO1 TM domain wild-type and mutants. **(B)** The fold-change in steady state levels of FLAG-HO1 wt and proline-266, arginine-269, leucine-272 and serine-275 residue mutants was examined by western blotting using anti-FLAG antibody. The quantification of the fold-change is shown below (mean ± SEM, n=3). **(C)** Influence of proline-266, arginine-269, leucine-272 and serine-275 residue mutants on FLAG-HO1 stability assessed by cycloheximide (CHX) chase and western blotting using anti-FLAG antibody (mean ± SEM, n=3) (see also Figure S6). **(D)** Steady-state western blot analysis of FLAG-HO1 wt, proline-266, arginine-269, leucine-272 and serine-275 mutants together with increasing concentrations of HA-SPP to detect SPP-generated cleavage fragments. Quantification shows reduced processing for the full-length form for the TM domain mutants. **(E)** Schematic representation of GFP-TEV-FLAG-HO1-TMD, its cleavage by SPP and further *in vitro* processing with TEV protease to generate small peptides for MS analysis. **(F)** Cleavage site mapping of HO1. The N-terminal fragment of the GFP-TEV-FLAG-HO1-TMD (as outlined in panel D) generated by ectopically expressed SPP was immunoprecipitated from the cytosol fraction after overnight treatment with 0.5 μM epoxomicin, digested with TEV protease and the resulting FLAG-tagged peptide was analyzed by MS. Cytosol from mock transfected cells was used as control. The inset shows high-resolution isotopic pattern of the 5274.76 Da peak.

In addition to the serine we showed that glutamine-197 in XBP1u plays a crucial role in SPP-dependent proteolysis. This residue is positioned four amino acids upstream of the serine-200 in XBP1u. In HO1, the corresponding residue four amino acids upstream of the critical serine-275 coincides with leucine-272. Although, glutamine and leucine have chemically different side chain properties they are similar in their spatial arrangement and may be important in positioning of the substrate into the catalytic active cleft. Thus, we mutated the leucine-272 to alanine and tested the effect in the cell based SPP assays. Notably, we observed that the L272A mutant of HO1 showed increased steady state levels (Figure 6B) and stability (Figure 6C). Taken together, these results indicate that a bipartite substrate feature including helix-destabilizing residues and a cleavage site motif is a common principle in the recognition of TM domain substrates by SPP.

For the HO1-S275A mutant a putative SPP-generated N-terminal cleavage fragment has been observed to localize to the nucleus in transfected HeLa cells (Hsu et al., 2015). Thus, we asked, to what extend these mutations prevent cleavage fragment generation. Because of a rapid proteasomal turnover, analyzing ectopically expressed HO1 we do not observe cleavage fragments generated by endogenous SPP, which is consistent with a previous report (Boname et al., 2014). Because of tight coupling to the ERAD machinery, fragments are commonly rapidly degraded by the proteasome (Boname et al., 2014; Chen et al., 2014). In fact, we previously observed that blocking the proteasome also suppressed SPP activity and only low amount of XBP1u cleavage fragments could be detected (Chen et al., 2014). The HO1 cleavage fragment, however, escapes complete proteasomal destruction (Hsu et al., 2015). To detect HO1 cleavage fragments without proteasome inhibition, we ectopically expressed increasing levels of SPP (Figure 6D) and compared processing of HO1 wt with the P266L, R269L, L272A and S275A mutants. Primarily, these results show that for all four mutants less substrate disappeared (Figure 6D), confirming their increased stability in presence of endogenous SPP levels (Figures 6B and C and S6A). However, at higher SPP levels also for these mutants an SPP-generated cleavage fragment was detected (Figure 6D). This indicates that upon overexpression, SPP may become more promiscuous and requirements of the cleavage site motif become less stringent recognizing alternative sites. Similarly, mutation of signal peptides can create alternative cryptic cleavage sites (Folz et al., 1988). However, taken together with the cycloheximide chase experiments, reduced processing efficiency of HO1 mutants on the endogenous protease level indicates that SPP engages in side chain-specific interactions with its substrates.

Next, we set out to determine the site of SPP-catalyzed cleavage. Previous analysis of SPP *in vitro* and tissue culture cells pointed towards cleavage in the middle of the TM span (Chen et al., 2014; Hussy et al., 1996; Lemberg and Martoglio, 2002; Weihofen et al., 2000). Because of tight coupling to the ERAD machinery with SPP-catalyzed XBP1u cleavage we failed to isolate sufficient XBP1u cleavage fragment for cleavage site determination (results not shown). HO1, however, escapes to some extent complete proteasomal destruction and a much less crosstalk of proteasome inhibition and SPP-trigged release has been observed (Boname et al., 2014; Hsu et al., 2015). Thus, making HO1 an ideal substrate for cleavage site determination by mass spectrometry (MS) analysis. To allow isolation of a short peptide comprising the carboxy-terminus generated by SPP, we co-expressed SPP with a HO1 TM domain GFP-fusion construct including an engineered TEV protease cleavage site followed by a FLAG-tag (Figure 6E). Subsequently, we immunoisolated the SPP-generated cleavage fragment from the cytosolic fraction with an anti-GFP antibody and then digested it on the beads with TEV protease. This leads to the release of 5 kDa-peptides containing the FLAG-tag at their new N-terminus and C-terminally ending at the peptide bond where SPP cleavage occurred. For cleavage site determination, these peptides were immunoprecipitated and analyzed by MS. A major peak was observed with 5274.76 Da, which corresponds to cleavage after threonine-273 (Figure 6F). In addition, three less intense peaks were observed corresponding to peptides further shortened by one amino acid, respectively. These alternate C-termini may either originate from SPP cleavage at alternative sites or from trimming by cytoplasmic carboxypeptidases. However, we did not detect any cleavage fragment matching the previously identified tryptic HO1 TM domain peptides of 6 to 7 residue length that indicated cleavage after serine-275 (Hsu et al., 2015). The predominant fragment observed by our analysis of 5-kDa peptides generated by a tailored TEV protease site strongly indicates that HO1 is predominantly cleaved at threonine-273. Taken together with the requirement for leucine-272 and serine-275, this suggest that SPP shows a so far unanticipated sequence preference at the P2 and P2’ position (according to Schechter and Berg’s nomenclature for protease subsites and substrate residues (Schechter and Berger, 1967)).

## Discussion

Here, we revisited XBP1u ER targeting and insertion into the ER membrane. We show *in vitro* that the XBP1u hydrophobic region can function as a signal peptide that productively interact with Sec61 leading to the subsequent translocation of a downstream protein domain. While previous work argues that XBP1u is primarily peripherally attached to ER-derived microsomes (Kanda et al., 2016; Plumb et al., 2015), by using a cell-based glycosylation reporter construct we show that at least half of the protein inserts as a type II membrane protein. Since glycosylation close to the C-terminal tail is commonly not complete, the insertion rate of XBP1u as a type II membrane protein may exceed 50%, verifying our previous topology studies including *in vitro* translocation assays with full-length XBP1u (Chen et al., 2014). Nevertheless, it cannot be ruled out that different ERAD pathways potentially also act on preinserted species and contribute to XBP1u degradation. Upon productive insertion as a type II membrane protein, XBP1u’s TM domain is metastable, prompting efficient recognition by SPP and intramembrane cleavage and subsequent proteasomal degradation. Mutagenesis studies in combination with amide DHX kinetics show that helix-destabilizing features are required for SPP-catalyzed cleavage. However, introducing a di-glycine motif in the C-terminal portion of the TM domain of XBP1u, which leads to local increase of the TM helix flexibility, does not interfere with ER localization but reduces SPP-triggered turnover. Site-specific mutagenesis of XBP1u and the TA protein HO1 shows that a serine residue and a long amino acid side chain promotes efficient SPP-catalyzed cleavage. Overall, our results suggest that SPP may recognize substrate sequence features like rhomboid intramembrane serine proteases (Strisovsky et al., 2009). However, our cleavage-site determination indicates that alternative cleavage sites exist. As observed for signal peptidase where multipe sites compete for cleavage (Folz et al., 1988), mutations of the substrate TM domain or overepxresison of SPP may generate new cleavage sites or lead to recognition of cryptic sites.

### Polar TM residues introduce functionality but affect ER targeting and helix stability

TM segments are 18 to 25 amino acid long a-helical peptides with a high content of hydrophobic residues. Their partitioning into the lipid bilayer of a cellular membrane via lateral exit from the Sec61 channel is thought to be driven by the hydrophobic effect (Cymer et al., 2014). While N-terminal and internal signal sequences are commonly targeted to Sec61 via SRP, thereby avoiding exposure of their hydrophobic domain to the aqueous environment, TA proteins are targeted and inserted into the ER membrane by the GET pathway (Hegde and Keenan, 2011). The thermodynamics of Sec61-mediated insertion of TM segments is described by ΔG_app_ determined by von Heijne and colleagues from measurements in a microsome-based system (Hessa et al.,
 2007). This scale reflects the situation of internal TM segments rather than that of moderately hydrophobic TM domains and N-terminal signal sequences (De Marothy and Elofsson, 2015; Dou et al., 2014; Junne and Spiess, 2017). Here, we show that in an isolated situation XBP1u’s TM domain is recognized by the Sec61 translocon leading to successful translocation (Figure 2B). Also, the C-terminus of ~50% of the steady state XBP1u pool is glycosylated, which is consistent with a predicted ΔG_app_ close to zero (Figure 2A). When this value decreases, as observed for several TM domain mutants investigated in this study (see Table S1), glycosylation efficiency increases up to >80% (Figures 1B and 3B). Likewise, the SPP-resistant XBP1u diglycine mutant (mt5) shows despite its higher flexibility in the DHX experiment (Figure 4C) a higher glycoslation rate, which is consistent with a decrease in its ΔG_app_ (Table S1). On the other hand, certain XBP1u mutants fail ER targeting. When fused to Prl as an N-terminal signal sequence, the XBP1u TM domain mt2, a mutationally rigidified TM helix variant, does not efficiently insert as a signal sequence. Consistent with this, N-terminal signal sequences frequently contain a helix-break-helix structure that is thought to favor loop-like insertion of the targeted nascent chain (van Klompenburg and de Kruijff, 1998). On the contrary, mt4, having the di-glycine motif in the center of the TM domain region, introduces helix flexibility and fails targeting. This is probably due to failed SRP interaction and/or ER insertion resulting from a positive ΔG_app_. These results show that for signal anchor sequences there is a fine balance between the number of hydrophilic residues that are tolerated and certain requirements for ER targeting and insertion. Despite that, polar amino acid residues in TM spans determine a plethora of important physiological functions. While we may not know the full set of XBP1u molecular properties and functions, here we show that its TM segment serves as a membrane-integral degradation signal that drives its turnover by an SPP-dependent ERAD pathway. Analogously, charged TM residues are recognized as so-called “degrons” by the ERAD machinery to target orphan receptor subunits for degradation (Bonifacino et al., 1990; Fleig et al., 2012).

### SPP-catalyzed cleavage is governed by conformational control and site-specific recognition

Since the first discovery of intramembrane proteolysis more than 20 years ago, an important unsolved question is how a lipid-embedded helical substrate is specifically recognized by a membrane-integral protease for cleavage (Brown et al., 2000; Langosch et al., 2015; Strisovsky, 2016; Wolfe, 2009). For several intramembrane protease substrates, a strict requirement for amino acid residues with a low predicted TM helix preference has been observed (Chen et al., 2014; Lemberg and Martoglio, 2002; Moin and Urban, 2012; Urban and Freeman, 2003; Ye et al., 2000). Therefore, it is commonly believed that unfolding of the substrate TM helix into the protease active site is a rate-limiting step (Lemberg and Martoglio, 2004). A flexible TM helix may be required to allow lateral movement of the scissile peptide bond from free lipid bilayer into the active site. For example, TM domain mutations in the amyloid precursor protein, a substrate of γ-secretase, can lead to Alzheimer’s disease and have an influence on processing. Such mutations have been proposed to change the global flexibility of the TM helix and thus to affect its positioning within the enzyme (Langosch et al., 2015; Stelzer et al., 2016). In addition, introducing glycines near the site where γ-secretase performs the first endoproteolytic cut, referred to as ε-site, facilitates cleavage (Fernandez et al., 2016) and solid-state NMR analysis revealed a helix-to-coil transition near the ε-site (Sato et al., 2009).

Here, we reveal an impact of the flexibility of the SPP substrate XBP1u TM helix by relating cell-based degradation assays to amide DHX kinetics. We observed that mutating hydrophilic residues either in the N- or C-terminal half of the TM domain to leucine locally stabilize the TM helix, which positively correlates with a failure in SPP-catalyzed turnover. Interestingly, upon introducing di-glycine motifs at the equivalent positions, we observed an increase in local helix dynamics as predicted, but no increase of cleavage efficiency. This is in line with our previous observation of flexibility requirements in the N-terminal TM domain portion (mt1) (Chen et al., 2014). However, the fact that the single-site mutation in S200G but not in Q199G mirrors the degradation kinetics seen with the C-terminal double-glycine mutant (mt5) implies that it is not only helix flexibility but also the primary structure determining the turnover efficiency by SPP. Likewise, our glycine scan revealed that glutamine-197 but no other tested residue is important for SPP-catalyzed turnover of XBP1u. Consistent with a model of site-specific substrate recognition, we observed that in the TA protein HO1 leucine-272 and serine-275, two positions upstream and two positions downstream of the scissile peptide bond, respectively, were both required for efficient cleavage by SPP. Hence, it is attractive to speculate that the SPP active site binds its substrates by the side chains of the P2 and P2’ positions, in a manner related to substrate discrimination by classical proteases (Schechter and Berger, 1967). Although the exact cleavage site is not known for the XBP1u TM domain, comparison with reference peptides suggests processing between residues 194 and 199 (Chen et al., 2014), which is in line with the model that also here a serine residue is placed in the P2’ position (see Figure 1 for XBP1u sequence). However, substrate recognition in P2 by of the other putative subsite may be less stringent tolerating glutamine-197 for XBP1u and leucine-272 for HO1. Likewise, slightly different results were observed with short signal peptides, where short h-regions completely lacking serine residues can be cleaved (Kronenberg-Versteeg et al., 2018; Lemberg and Martoglio, 2002). This suggests that there is no strict need for a particular side chain, but rather a combination of different factors accelerate SPP-catalyzed intramembrane cleavage. However, we showed that the neo-signal peptide derived from the XBP1u TM domain is not efficiently cleaved when the serine is replaced by a di-glycine motif (Figure 3E), highlighting the relevance of this feature also for signal peptide processing. Based on our results, we propose a two-step model for substrate recognition by SPP (Figure 7). First, the helical substrate TM segment is recognized by a substrate-binding site on SPP that is remote from the active site, referred to as “exosite”, as it has been suggested for other intramembrane proteases (Arutyunova et al., 2014; Fukumori and Steiner, 2016; Kornilova et al., 2005; Strisovsky et al., 2009). We postulate that in a second step the substrate TM helix subsequently partially unwinds with the peptide comprising the scissile peptide backbone binding into a putative active site grove. Consistent with this, a recent study applying resonance Raman spectroscopy revealed unwinding of the substrate helix into a β-strand geometry upon binding to an archaeal presenilin/SPP homologue (PSH) (Brown et al., 2018). Prior to local unwinding for cleavage, flexibility in the N-terminal half of the TM helix may be required for substrate translocation and its correct positioning (Fig 7A). Analogously, h-regions of signal peptides may have an intrinsic tendency to enter the SPP active site to minimize the energetically disfavored hydrophobic mismatch caused by their short span through the lipid bilayer (Figure 7B). While for short substrates this may be the only rate-limiting determinant, for more stable TM helices the equilibrium may be more on the side of the helical conformation and only a cognate recognition motif tightly docking into the active site may be efficiently bound and cleaved (Fig. 7A). Likewise, recent kinetic measurements using the purified archaeal PSH showed a preference for cleavage at a threonine at the scissile peptide bond (Naing et al., 2018). Although the mechanism of substrate selection may be different, in an intriguing parallel also rhomboid proteases combine recognition of a defined cleavage site motif (Strisovsky et al., 2009) with sampling TM domain dynamics (Akiyama and Maegawa, 2007; Moin and Urban, 2012; Strisovsky, 2016; Urban and Freeman, 2003). Overall, as known from limited proteolysis by soluble proteases, the emerging picture is that also in the lipid bilayer a continuum of conformational control and site-specific substrate recognition governs protease selectivity. We note, however, that this model remains speculative until the structure of an SPP-type protease with its substrate has been determined.

**Figure 7.**
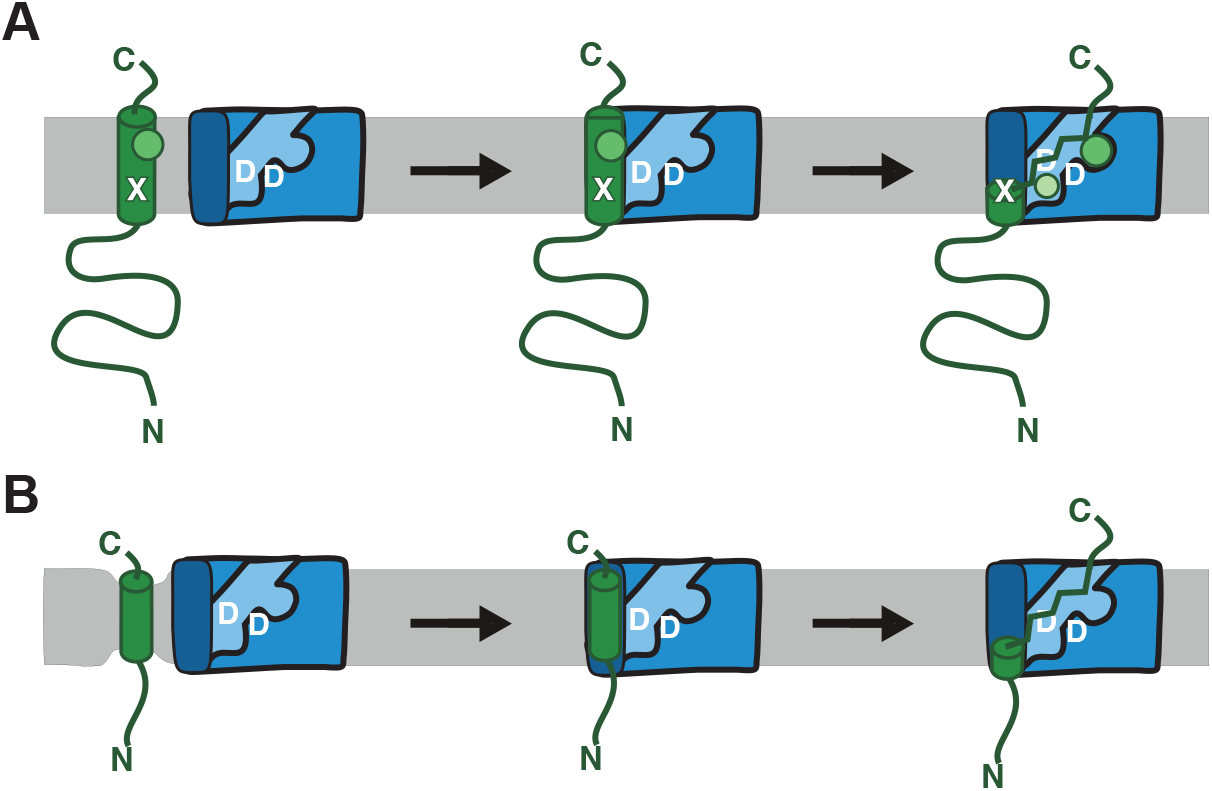
Model for SPP substrate recognition. **(A)** SPP binds helical TM spans of type II membrane proteins and TA proteins by a putative docking site (dark blue), which despite its predicted low affinity may serve as an “exosite” introducing the first layer of specificity. In order to expose the scissile peptide bond to the active site (bright blue, “D” indicates catalytic residues), substrate TM helix has to partially unfold, a reaction that is facilitated by helix-destabilizing features (X). This equilibrium may be shifted to the open conformation by site-specific recognition of substrate features such as serine (green circle) and a long amino acid side chain (light green circle) that are bound to the active site cleft via putative “subsites”. Reaction rate may be determined by helix-dynamics (rate controlled) and the primary sequence (affinity driven). **(B)** With an h-region in the range of 7 to 15 amino acids, TM-spans of signal peptides are commonly too short to form a regular TM helix and may directly unfold into the SPP active site to minimize the high energetic state of hydrophobic mismatch in the surrounding lipid bilayer. Subsequently, side-specific contacts to a cleavage site region is not a strict requirement and cleavage is only rate controlled.

## Experimental Procedures

### KEY RESOURCES TABLE

See separate excel file.

## CONTACT FOR REAGENT AND RESOURCE SHARING

Further information and requests for reagents should be directed to and will be fulfilled by the Lead Contact, Marius K. Lemberg.

## EXPERIMENTAL MODEL AND SUBJECT DETAILS

### *E. coli* strains XL-10 Gold

*E. coli* strain XL-10 Gold (Stratagene) was used for DNA plasmid amplifications. The strain was grown in LB-medium or on LB-agar with when transformed depending on the resistance gene of the corresponding plasmid was either ampicillin or kanamycin.

### Cell lines and transfection

Hek293T cells were grown in DMEM (Invitrogen) supplemented with 10% fetal bovine serum (FBS) at 37°C in 5% CO_2_. Transient transfections were performed using 25 kDa linear polyethylenimine (Polysciences) (Durocher et al., 2002). Typically, 1 μg of XBP1u plasmid and 300 ng of HO1 plasmid were used per well of a 6-well plate (see supplemental information for details on the plasmids). Total transfected DNA (2 μg/well) was held constant by the addition of empty plasmid. If not otherwise stated, cells were harvested 24 hours after transfection. To inhibit the proteasome cells were treated with 1 μM epoxomicin (Calbiochem); to inhibit SPP, 50 μM (Z-LL)_2_-ketone (Calbiochem) in dimethyl sulfoxide (DMSO) was added for 16 hours and subsequently harvested for western blot analysis.

## METHOD DETAILS

### Cloning of Plasmids

For the cell-based assays, if not otherwise stated, all open reading frames were cloned into pcDNA3.1+ (Invitrogen). Plasmids encoding human XBP1u (harboring a silent mutation in the 3’-IRE1-splice-site) tagged with an N-terminal triple FLAG-tag and SPP with triple HA-tag inserted between residue 373 and the C-terminal KKXX ER-retention signal (SPP-HA) had been described previously (Chen et al., 2014). The XBP1u TM domain mutants mt1 (S187L, P188L, A192L), mt2 (Q197L, Q199L, S200L, S203L), mt3 (S187G, P188G), mt4 (L194G, T195G), mt5 (Q199G, S200S), L196G, Q197G, I198G, Q199G, S200G, L201G, I202G, S203G, C204G as well as the R232N glycosylation mutants were generated by Quick-Change site-directed mutagenesis (Stratagene). The deletion constructs i.e. the tail-anchored XBP1u mutants were generated by subcloning the respective region of the open reading frame (1-207). HO1 (full-length ORF Gateway clone 187935150) was cloned into pcDNA3.1 with an N-terminal triple FLAG-tag and the point mutations (P266L, R269L, L272A and S275A) were introduced by Quick-Change mutagenesis. For the *in vitro* experiments all constructs were cloned into pRK5. The construct coding bovine pPrl has been described previously (Schrul et al., 2010). The XBP1u-TMD-Prl, wt, mt1 and mt2 were ordered as gene blocks (IDT) comprising amino acid sequence 177-207 of XBP1u (wt or the respective mt) followed by the mature Prl sequence (31-229). XBP1u-TMD-Prl mt4 was generated by Quick-Change site-directed mutagenesis (Stratagene) by using XBP1u-TMD-Prl wt as a template. In order to map the SPP-cleavage site in transfected cells, the open reading frame encoding residues 237-288 of HO1 was cloned into pEGFP-C1 (Invitrogen) proceeded by a linker contacting a TEV protease cleavage site and a single FLAG-tag (…ENLYFQ^GDYKDDDDKG…). The ER marker ssRFP-KDEL were previously described (Snapp et al., 2006).

### Cycloheximide chase experiments

Cycloheximide (100 μg/ml) chase was conducted 24 hours after transfection of Hek293T cells, and cell extracts were subjected to western blot analysis. To inhibit SPP, 50 μM (Z-LL)_2_-ketone (Calbiochem) in dimethyl sulfoxide (DMSO) was added 2 hours before the addition of cycloheximide and the inhibitor was kept on the cells during the chase experiment.

### SDS-PAGE and western blot analysis

Proteins were solubilized in SDS sample buffer (50 mM Tris-Cl pH 6.8; 10 mM EDTA, 5% glycerol, 2% SDS, 0.01% bromophenol blue) containing 5% β-mercaptoethanol. All samples were incubated for 15 min at 65°C. For deglycosylation, SDS sample buffer solubilized proteins were diluted 1:4 in water to reduce the SDS contentration to 0.5% and were treated with EndoH (New England Biolabs) according to the manufacturer’s protocol. For western blot analysis, proteins were separated by SDS-PAGE using Tris-glycine acrylamide gels and transferred to PVDF membrane followed by enhanced chemiluminescence analysis (Pierce) to detect the bound antibodies. The following primary antibodies were used: mouse monoclonal anti-FLAG (M2, Sigma; 1:1000), mouse monoclonal anti-HA (BioLegend; 1:1000) mouse monoclonal anti-β-Actin (Sigma; 1:4000). The following secondary antibodies were used: Donkey-anti-mouse IgG–HRP conjugated (Dianova; 1:10000). For detection the LAS-4000 system (Fuji) was used. Data shown are representative of three independent experiments. For quantification we used FIJI (Schindelin et al., 2012) and data acquired from the LAS-4000.

### Microscopy

For immunofluorescence detection of N-terminally FLAG-tagged XBP1u, transfected Hek293T cells were grown on glass cover slides and fixed in PBS containing 4% paraformaldehyde for 30 min. Permeabilization was performed in PBS containing 0. 5% Triton X-100 for 10 min. After blocking cells for 30 min in PBS containing 20% FBS and 0.5% Tween-20, cells were incubated in PBS/FBS/Tween-20 with the primary antibody, mouse monoclonal anti-FLAG (M2, Sigma; 1:1000), for 1 h at room temperature. Subsequently, cells were washed in PBS/FBS/Tween-20, incubated with fluorescent secondary antibody, Donkey-anti-mouse IgG Alexa Fluor 488 (Invitrogen; IF 1:2000), washed, stained with Hoechst 33342 (Thermo Fisher Scientific) and mounted for analysis by confocal microscopy. For selective permeabilization of PFA-fixed cells on a glass cover confocal microscopy was performed on a Zeiss LSM confocal microscope. Images were taken with a Plan-APOCHROMAT 63×/1.4 oil objective lens with a pinhole setting of 1.0 Airy unit. Image processing was performed using FIJI (Schindelin et al., 2012).

### *In vitro* transcription, translation/translocation, protease protection assay and signal peptide processing

pRK5-Prl or pRK5-XBP1uTMD-Prl plasmids were linearized with XhoI and transcribed with the SP6 RNA polymerase at 42°C in the presence of 500 μM m7G(5’)ppp(5’)G CAP analogue (New England Biolabs). mRNA was translated in rabbit reticulocyte lysate (Promega) containing [^35^S]-methionine/cysteine as described before (Lemberg and Martoglio, 2002). Where indicated, nuclease-treated rough microsomes prepared from dog pancreas were added (Martoglio et al., 1998). For protease protection, the reactions were treated with 0.5 mg/ml proteinase K for 45 min on ice. As a control, proteinase K was added in the presence of 1% Triton X-100. Subsequently, 10 μg/ml PMSF was added, and samples were analyzed on 12% Tris-glycine acrylamide gels and autoradiography as described previously (Lemberg and Martoglio, 2002). To inhibit SPP processing in the signal peptide processing assay 10 μM (Z-LL)2-ketone was added. Samples were incubated 20 min at 30°C. Microsomes were extracted with 500 mM KOAc and analyzed on Tris–Bicine acrylamide gels (15% T, 5% C, 8 M urea). Signal peptide processing was quantified by comparing intensity of signal peptide in the absence (DMSO) and presence of SPP inhibitor (Z-LL)2-ketone for each condition, given by the formula 100-(100x/y) where x is the intensity of the test band and y is the intensity of the band in the presence of full inhibition with (Z-LL)2-ketone. Equal translocation efficiency was controlled by comparing the amount of Prl between conditions. Translation of the reference peptide containing the XBP1u-TMD was performed in wheat germ extract at 25°C for 30 min (Martoglio et al., 1998).

### Mass spectrometry-based mapping of SPP-cleavage site

Hek293T cells were transiently transfected with GFP-TEV-FLAG-HO1-TMD fusion construct and SPP-HA and treated 24 hours post transfection with 0.5 μM epoxomicin for 16 h. Subsequently, cells were detached in cold PBS-EDTA and resuspended in hypotonic buffer (10 mM HEPES-KOH pH 7.4, 1.5 mM MgCl_2_, 10 mM KCl, 0.5 mM DTT, 10 μg/ml phenylmethylsulfonyl fluoride (PMSF), and each of 10 μg/ml chymostatin, leupeptin, antipain and pepstatin). After 10 min incubation on ice, cells were lysed by passing five times through a 27-gauge needle. Cell lysates were cleared by centrifugation at 5,000 g for 5 min and 20,000 g for 30 min at 4°C and the obtained cytosolic fraction was pre-absorbed for 1 h on BSA-coupled sepharose beads. Anti-GFP immunoprecipitation was performed for 30 min at room temperature using a GFP-specific single chain antibody fragment coupled to sepharose beads as described (Fleig et al., 2012). Immunoprecipitates were washed three times in buffer WB (50 mM HEPES-KOH, pH 7.4, 150 mM NaCl, 2 mM MgOAc2, 10% glycerol, 1 mM EGTA, 0.1% Triton X-100) followed by washing once in buffer TDB (50 mM Tris-HCl, pH8.0, 0.5 mM EDTA, 1 mM DTT, 5 μg/ml PMSF, 10% glycerol, 0.1% Triton X-100). Subsequently immunoprecipitated proteins were digested by adding 3 μl recombinant TEV protease to the beads suspended in buffer TDB overnight at 20°C on a rotary shaker. Supernatant and buffer from three wash steps with PBS were pooled and eluted GFP-tagged peptides were first immunoabsorbed with GFP-specific single chain antibody beads, followed by immunoprecipitation of FLAG-tagged peptides with FLAG M2 agarose (Sigma). Agarose was washed two times with PBS followed by two times with ddH_2_0. Peptides were eluted from the beads with 15 μl 1% TFA/10% acetonitrile. 5μl eluate was analyzed by a nanoHPLC system (Ultimate RSLC 3000) coupled to an ESI QExactive mass spectrometer (Thermo Fisher). The sample was loaded on a self-packed analytical column (75um x 250mm, Reprosil-Pur C-18-AQ 1.9um (Maisch GmbH) and eluted with a flow rate of 300nl/min in a 60 min gradient of 3% to 40% buffer B (0.1% formic acid, acetonitrile). Buffer A was 0.1% formic acid. One survey scan (res: 120.000) was followed by 10 information dependent product ion scans.

### *In silico* TM domain analysis

To compare the XBP1u TM region, signal sequences and canonical TM segments we determined the free energy differences (ΔG_app_) by the online ΔG prediction server (http://dgpred.cbr.su.se). To predict the presence and the site of signal peptide cleavage we used SignalP 3.0 (http://www.cbs.dtu.dk/services/SignalP-3.0) and TM domain mutants were evaluated using TMHMM (http://www.cbs.dtu.dk/services/TMHMM).

### Peptide synthesis

Peptides were synthesized by Fmoc chemistry (PSL, Heidelberg, Germany) and purified by HPLC. They were > 90% pure as judged by mass spectrometry. Concentrations were determined via UV spectroscopy using an extinction coefficient at 280 nm of 11 000 M^-1^cm^-1^.

### Circular dichroism spectroscopy

For CD spectroscopy, peptides were dissolved at 50 μM in 80% (v/v) TFE, 20% (v/v) 10 mM Tris/HCl (respectively 5 mM PBS for mt5 and mt3), pH 7. CD spectra were obtained using a Jasco J-710 CD spectrometer from 190 nm to 260 nm in a 1.0 mm quartz cuvette at 20°C with a response of 1 s, a scan speed of 100 nm/min and a sensitivity of 100 mdeg/cm. Spectra were the signal-averaged accumulations of 10 scans with the baselines (corresponding to solvent) subtracted. Mean molar ellipticities were calculated and secondary structure contents estimated by deconvoluting the spectra using CDNN software (Bohm et al., 1992) with CDNN complex reference spectra.

### Mass spectrometry and DHX experiments

Peptides were dissolved in 80% (v/v) d1-HFIP/20% D2O at a concentration of 300 μM and incubated at 37°C for 7 days, solvent was removed by vacuum centrifugation, and the pellet was re-dissolved in deuterated solvent (80% (v/v) d1-TFE, 2 mM ND4-acetate). This resulted in > 95% deuteration as determined by ESI-MS. Deuterated DTT was prepared by repeatedly dissolving DTT in D2O and lyophilizing overnight; the deuterated DTT was finally dissolved in D2O. Deuterated DTT was added at 1 mM to prevent cysteine oxidation in the deuterated peptides. For measurements of overall DHX, the deuterated peptides were diluted 1:20 from a 100 μM stock solution into 80% (v/v) TFE, 2 mM NH4-acetate, 1 mM tris(2-carboxyethyl)phosphine (TCEP), pH 5.0, and incubated for different time periods at 20.0°C after which DHX was quenched on ice and by adding (0.5% (v/v) formic acid (pH ≈2.5). For electron transfer dissociation (ETD), 300 μM stock solutions were diluted 1:20. Mass/charge (m/z) ratios were recorded after the indicated time periods by electron-spray-ionization mass spectrometry (ESI-MS) as described (Poschner et al., 2009) using a Synapt G2 HDMS (Waters Co.) with one scan/second and evaluated as described (Stelzer et al., 2008). For ETD-DHX, incubation times from 1 min to 3 days at pH 5.0 were chosen. An incubation time of 0.1 min at pH 5 was simulated by incubation for 1 min at pH 4 and an incubation time of 30 days at pH 5 was simulated by incubating for 22 hour at pH 6.5. ETD was conducted as described (Stelzer et al., 2016) at a flow rate of 3 μl/min using a cooled syringe. ETD was started by decreasing the wave height from 1.5 V to 0.2 V. Hydrogen scrambling was < 5-10%, as determined by the ammonia loss method (Rand et al., 2012). All experiments were done at least in triplicate.

ETD fragment spectra recorded over 10 min were combined and evaluated using the module *ETD Fragment Analyzer* of the software suite MassMap^®^ (MassMap GmbH & Co. KG, Freising, Germany) that is based on GRAMS/AI (Thermo Fisher Scientific). The theory and the detailed evaluation procedure are described in the section *“Analysis of amide-exchange/ETD Data using the ETD Fragment Analyzer*” (see below). Briefly, the evaluation consisted of the following steps:

1. Determination of the c- and z-fragment ions to be included in the analysis as well as of the ‘extra hydrogens’ of the fragment ions from the ETD spectrum of a non-deuterated sample. ‘Extra hydrogens’ result from hydrogen transfer during fragmentation or from neutralization of precursor ion proton by electron transfer (Lermyte et al., 2015).
2. After DHX and ETD, the expected isotopic patterns of the fragment ions (determined in step 1) were calculated by considering the rapid exchange of labile deuterons linked to side-chain heteroatoms or the N- and C-terminus, charge carrying deuterons, and extra deuterons (determined in step 1).
3. The calculated isotopic patterns were compared to the experimental ones by determining the extents of amide deuteration.
4. To increase the precision, a smoothing and interpolation procedure was applied to the mean numbers of deuterons as a function of the fragment numbers by fitting second order polynomials to four consecutive data points. The extent of amide deuteration was then determined from the smoothed and interpolated mean deuteron numbers of fragment ions.
5. The numbers of amide deuterons as a function of exchange period D(t) as obtained in step 4) were used to compute the residue-specific first order exchange rate constant k_exp_ using Eq. (1) (which considers 5% remaining deuterated solvent)

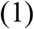
6. The equilibrium constants K_exp_ were calculated from K_exp_ and the sequence specific chemical exchange rate constants k_ch_ based on Eq. (2) based on Linderstrom-Lang theory, assuming EX2 conditions and a predominant folded state (Skinner et al., 2012)

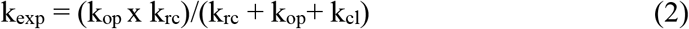

where k_op_ and k_cl_ are the rate constants of H-bond opening and closing, respectively, and krc represents the chemical rate constant that was calculated by http://landing.foxchase.org/research/labs/roder/sphere/ (under the set conditions: D-to-H-exchange, reduced Cys, pH =5.0, T = 20.0°C).
7. The free energies ΔG required for H-bond opening were then calculated according to Eq. (3)

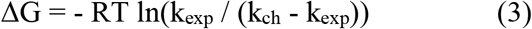 It should be noted, that the ΔΔG values obtained with this procedure are an upper estimate of the true values since (i) the molarity of water in 80% (v/v) TFE solvent is only 20% of the bulk molarity used for the determination of the reference chemical exchange rates k_ch_, and (ii) the hydration of residues in the hydrophobic core of a TMD is possibly reduced relative to bulk. Both factors likely reduce the chemical exchange rate in our experiments. In addition, TFE might have an impact on the autoionization constant of water and the chemical exchange rate constants (Stelzer et al., 2016).

### Analysis of amide-exchange/ETD Data using the ETD fragment analyzer

#### 1. Definition of Terms and Abbreviations

| **Term** | **Meaning** |
| --- | --- |
| Asymptotic D number | Relative deuteron abundance of exchanging hydrogens that is asymptotically approached for very long exchange periods. The limit equals the ratio of deuterated to hydrogenated solutions employed in exchange. |
| H/D exchange | Experiment where H is exchanged by D |
| D/H exchange | Experiment where D is exchanged by H |
| D number | For a fragment ion: mean number of deuterons occupying the locations of slowly exchanging amide hydrogens<br>For an amide hydrogen: relative deuteron abundance of the amide hydrogen |
| Extra hydrogens | Hydrogens that either result from the transfer of hydrogens as part of the fragmentation process or result from the neutralization of charge carrying hydrogens of the precursor ions by means of electron transfer |
| Fast exchange | Exchange observed for exchangeable hydrogens other than amide hydrogens. Fast exchange is observed for four types of hydrogens: hydrogens linked to heteroatoms of the amino acid side chains, hydrogens linked to the heteroatoms of the functional groups of the N and C terminus, charge carrying hydrogens, extra hydrogens (for definition see above). |
| Isotopic number | Number counting the isotopic peaks of an isotopic cluster starting with 0 for the monoisotopic peak. The number is incremented by 1 for the addition of a single nominal mass unit ($^{12}\text{C}$ replaced by $^{13}\text{C}$, addition of a single extra hydrogen, $^1\text{H}$ replaced by $^2\text{H}$). |
| Slow exchange | Exchange observed for amide hydrogens and slow enough to be followed experimentally |

| **Abbreviation** | **Meaning** |
| --- | --- |
| FWHM | Full width at half maximum height/intensity |
| i | Isotopic number |
| i<sub>max</sub> | Maximum isotopic number considered for fragments including extra hydrogens |
| i<sub>limit</sub> | Maximum isotopic number considered for the calculated isotopic pattern of a neutral fragment without extra hydrogens. The limitation of the isotopic numbers is employed in order to avoid the consideration of isotopic peaks whose intensity is too low to be of practical relevance. |
| I | Expected isotopic distribution of a neutral fragment without extra hydrogens |
| I<sub>xxx</sub>, I<sup>(k)</sup> | Different isotopic distributions employed for the analysis of experimental isotopic patterns (xxx = +j, fit, ...; k = 1,2) |
| m<sub>av</sub> | Average mass of a neutral fragment without extra hydrogens |
| m<sub>mi</sub> | Monoisotopic mass of a neutral fragment without extra hydrogens |
| z | Charge number of a fragment |
| z<sub>precursor</sub> | Charge number of the precursor ion |

#### 2. Definition of an amide-exchange/ETD project

The pieces of information and the data needed for an amide-exchange/ETD project are as follows:

- Information sufficient for the calculation of the expected full width at half maximum intensity (FWHM) of isotopic peaks (type of mass analyzer, resolution function).
- Amino acid sequence of the peptide.
- If applicable, status of Cysteine residues (Cysteine residues reduced or alkylated by means of one of the common alkylation reagents).
- For all modified amino acids, positions and type of modification, e.g. acetylation of the N-terminus or amidation of the C-terminus.
- Type of the exchange experiment, i.e. D exchanged by H (below referred to as D/H exchange) or H exchanged by D (H/D exchange).
- Asymptotic D number depending on the ratio of deuterated to hydrogenated solutions employed in exchange.

In the case of a D/H exchange experiment, a fully (>95%) deuterated peptide dissolved in a fully deuterated medium (‘starting solution’) is mixed with a fully hydrogenated solution (‘dilution solution’). If the volume of the starting solution and the volume of the dilution solution are abbreviated by VD and VH, respectively, the asymptotic D number is calculated as the quotient V_D_/(V_D_+V_H_). For H/D exchange, the asymptotic D number is calculated in exactly the same way. The only difference is the meaning of the two volume variables: V_H_ and V_D_ then stand for the volume of the starting solution (fully hydrogenated medium containing the peptide) and the volume of the dilution solution (fully deuterated medium without the peptide), respectively.

The term ‘asymptotic D number’ is based on the expectation that the degree of deuteration of the slowly exchanging amide hydrogens will asymptotically approach that number for very long exchange times. For the calculations of the isotopic distributions of the peptide and fragment ion clusters, it is assumed that the degree of deuteration of all fast exchanging hydrogens equals the asymptotic D number. The term ‘fast exchanging hydrogens’ subsumes all the non-amide exchanging hydrogens, e.g. the side chain hydrogens linked to heteroatoms. In the formulas shown below, the asymptotic D number is abbreviated by D_asymptotic_.

- Charge number of the precursor ion (z_precursor_).
- Data sets with fragment scans acquired for different samples under exactly the same conditions, i.e. with exactly the same solvents and gases as well as with the same parameters of the ion source and the mass analyzer.

As part of the evaluation, sum mass spectra are calculated using either all the scans or the scans of a user-selected scan interval.

One of the samples, below referred to as ‘non-deuterated’, is the fully hydrogenated peptide dissolved in the fully hydrogenated medium. The other samples (‘exchange samples’) are mixtures of the starting solution and the dilution solution. The mixtures only differ by the exchange period, i.e. the length of the time interval ranging from the time of mixing to the time of mass spectrometric analysis.

#### 3. Evaluation of amide-exchange/ETD Project

The evaluation is implemented as a particular module of the MassMap^®^ software suite called *ETD Fragment Analyzer*. The module was designed in order to perform as many tasks as possible in an automated manner and, at the same time, to allow for full user control of all relevant decisions.

The evaluation of a single amide-exchange/ETD project consists of several steps that are described in the following sections.

##### 3.1. Evaluation of the non-deuterated sample

The first step is the evaluation of the sum mass spectrum of the non-deuterated sample. It is the aim of this step to identify the fragment clusters and to determine the number of neutral hydrogens of the fragments. Neutral hydrogens either result from the transfer of hydrogens as part of the fragmentation process or result from the neutralization of charge carrying protons of the precursor ions by means of electron transfer. These neutral hydrogens will be referred to as ‘extra hydrogens’ below. The very first decision is made by the user who defines the types of fragment ions (x, x*, x° with x=a, b, c, x, z, y; * and °: ions with neutral loss of ammonia or water, respectively) to be investigated. For the charge numbers (z) that are also selected by the user, the program automatically analyzes the isotopic patterns of all possible fragment clusters. The analysis of the isotopic patterns is based on the count sums over the extraction intervals of the isotopic peaks. The isotopic numbers (i) of the peaks range from 0 (isotopic number of the monoisotopic peak of the fragment without extra hydrogen) to i_max_. To determine i_max_, the normalized isotopic pattern or distribution I of the fragment is approximated by a binomial distribution characterized by the effective number n_C_ of carbon atoms. The number n_C_ is calculated by means of the monoisotopic mass m_mi_ and the average mass m_av_ of the fragment in a way that n_C_ · (mc_13_ – mc_12_) · a_13_ equals the difference m_av_ – m_mi_. The quantities mc_13_, mc_12_ and a_13_ stand for the mass of ^13^C, the mass of ^12^C and the natural abundance of ^13^C amounting to 0.011, respectively. Hence, the elements I(i) of the isotopic distribution I are calculated according to equation (1):

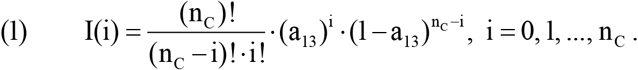

Only the peaks with isotopic numbers ranging from 0 to i_limit_ are considered. The upper limit of the isotopic numbers i_limit_ is the smallest natural number meeting the following criterion:

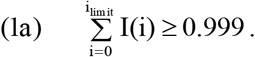

In order to take the extra hydrogens into account that are caused by the fragmentation process itself or by neutralized charge carrying protons of the precursor ion, the largest isotopic number i_max_ for which the count sum is calculated is determined by equation (1b):

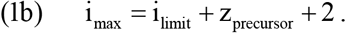

The limits mz_lower, i_ and m_zupper, i_ of the extraction interval of an isotopic peak with isotopic number i is based on the FWHM of the isotopic peak defined by the settings of the mass analyzer. In the following, the center m/z of the isotopic peak is termed mzi. The quantities mz_i_, mz_lower, i_ and mz_upper, i_ are calculated in the following way:

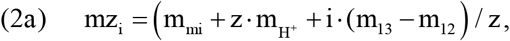

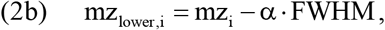

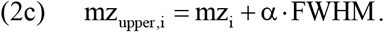

By default, the value of factor α amounts to 1. The value may be slightly changed by the user in order to avoid cross-talk from interfering peaks or in order to make sure that all the relevant counts of the isotopic peaks of the clusters are taken into account.

The count sum C(i) of the isotopic peak with isotopic number i is calculated as the sum of the intensities of the data points of the sum mass spectrum of the non-deuterated sample, whose m/z values are within the interval [mz_lower, i_; mz_upper, i_].

In cases of a calibration error of the mass spectrometer, the user may select two peaks of the sum mass spectrum that are used for a linear recalibration of the m/z axis of the mass spectrum. The recalibration option is not restricted to the mass spectrum of the non-deuterated sample. Rather, it is available for all the sum mass spectra of an amide-exchange/ETD project.

As the last step of the evaluation of fragment clusters of the non-deuterated sample, multiple linear regression is performed in order to determine the distribution of extra hydrogens of the clusters. For that purpose, the contributions of the unchanged isotopic distribution I as calculated by equation (1) (corresponding to fragment ions without extra hydrogens) and of the shifted isotopic distributions I+j with j=1, 2,…, z_precursor_+2 are determined. The shifted distribution I_+j_ corresponds to the expected distribution of the fragment cluster with j extra hydrogens:

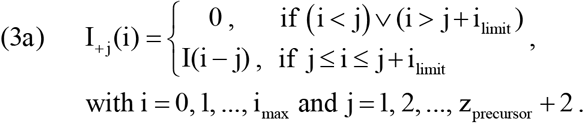

To determine the contributions of the different distributions to the experimental distribution C (composed of the count sums C(i) with i=0, 1,…, i_max_), the coefficients cj (j=0, 1,…, y_precursor_+2; I+0 = I) and the normalization factor a are calculated by minimizing the quantity χ^2^ of equation (3b):

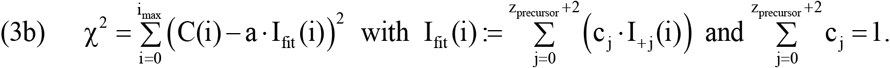

I_fit_ denotes the normalized isotopic distribution fitted to the experimental distribution. The fitting procedure defined by equations (3a) and (3b) is performed automatically as soon as a cluster is activated. In order to eliminate mass spectrometric peaks that are most probably part of other clusters, peaks with mass deviations above a user-defined limit or with too high or too low intensities are excluded from the fit by means of an iterative procedure. The final decision, which peaks are to be included in the fit, is made by the user. In most cases, intense clusters well above the limit of detection do not need user action, whereas for clusters around that limit user action is frequently needed in order to get valid results.

##### 3.2. Evaluation of the exchange samples

The second step is the evaluation of the sum mass spectra of the samples having undergone exchange. In principle, the procedure applied is the same as the procedure employed for the analysis of the clusters of the sum mass spectrum of the non-deuterated sample. The differences between spectra with and without prior exchange are due to the fact that quickly exchanging hydrogens as well as slowly exchanging hydrogens have to be taken into account. Fast exchange, i.e. exchange too fast to be followed experimentally, is to be considered for the extra hydrogens (for definition see the previous section), for the side chain hydrogens linked to heteroatoms, for the carboxyl and the amino hydrogens located at the N or C termini as well as for the charge carrying protons. The slowly exchanging hydrogens are non-hydrogen bonded amide hydrogens of the peptide backbone. The isotopic distribution of a cluster of an exchange sample is calculated from the isotopic distribution I_fit_ of the respective cluster of the non-deuterated sample by a series of successive convolution steps.

The first convolution steps are due to the fast exchange of the extra hydrogens. The extra hydrogens of the ion species with extra hydrogens are assumed to be fast exchanging. For these hydrogens, the probability for the incorporation of a D instead of an H is assumed to be equal to the asymptotic D number Dasymptotic. Hence, the additional distribution to be considered for the ion species with j extra hydrogens is a binomial distribution denoted by I_extra, j_:

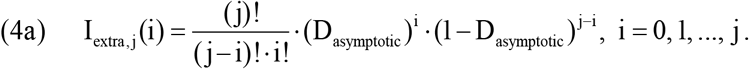

As a consequence, the fast exchanging extra hydrogens will change the distribution Ifit into the distribution I^(1)^ that is calculated in the following way:

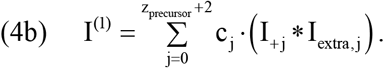

The notation f*g stands for the convolution of the two discrete distributions f and g. If f and g have n_f_+1 and n_g_+1elements with indices ranging from 0 to n_f_ and n_g_, respectively, h=f*g constitutes a distribution with the (n_f_ + n_g_ +1) elements h(i):

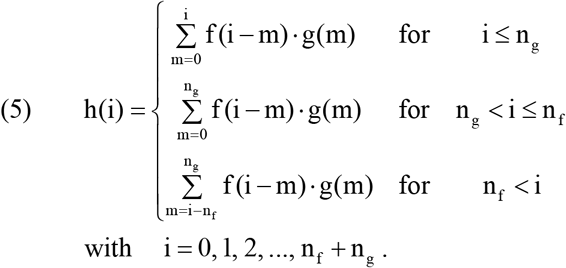

In addition to the extra hydrogens that change the expected isotopic distribution from I_fit_ to I^(1)^, the fast exchanging charge carrying protons and the fast exchanging hydrogens covalently linked to heteroatoms have to be considered.

If the number of these hydrogens is denoted by n_fast_plus_, the distribution I^(2)^, i.e. the distribution expected in the case of lacking slow exchange, is the convolution of the binomial distribution I_fast_plus_ of the n_fast_plus_ fast exchanging hydrogens with the distribution I^(1)^ of equation (4b):

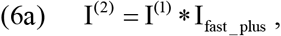

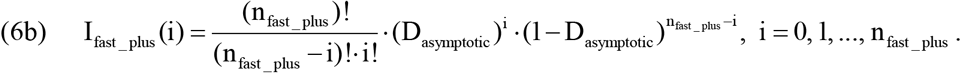

The final distribution that is to be compared with the experimental distribution C found in the mass spectrum of the exchange sample is the convolution of distribution I^(2)^ with the distribution of the slowly exchanging hydrogens.

From experience, for a significant fraction of the experimental clusters, satisfying fits are only obtained by employing a superposition of two binomial distributions, i.e. a bimodal binomial distribution for the distribution of the nslow slowly exchanging hydrogens. Apart from nslow, such a bimodal distribution is characterized by three numbers, namely the fraction fα of the first of the partial distributions and the exchange probabilities α and β of the first and the second partial distribution, respectively. Thus, the calculation of the distribution Islow of the nslow slowly exchanging hydrogens is performed in the following way:

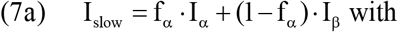

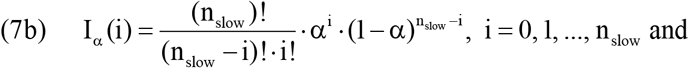

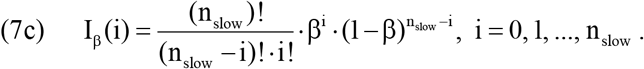

The convolution of the distribution I^(2)^ and the distribution Islow results in the distribution I^(3)^ that is fitted to the distribution C made up by the count sums of the isotopic peaks of the fragment cluster of interest. The numbers f_α_, α and β, all ranging between 0 and 1, as well as a normalization factor (defined as in equation (3b)) are obtained by a non-linear least squares fitting procedure. The mean number of deuterons D_cluster_ occupying the positions of the slowly exchanging amide hydrogens of the cluster is calculated according to the following equation:

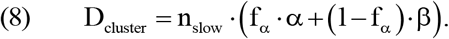

##### 3.3. Determination of the D numbers of the slowly exchanging amide hydrogens

The third step consists in the determination of the D numbers of individual slowly exchanging amide hydrogens. For the calculation of the D number D_k_ of the amide hydrogen of amino acid residue k (k=1, 2,…,n_A_S; n_A_S: number of amino acids of the peptide), suitable pairs of consecutive fragment ions of the same type are used. To be suitable for analysis, a given fragment ion (A) must contain a single additional amide hydrogen relative to another fragment ion (B). The D number Dk of that amide hydrogen is calculated as the difference D_A_ – D_B_.

As an option, the D numbers of the clusters of a particular type of fragments can be smoothed and interpolated before they are used for the determination of the D numbers of the individual amide hydrogens. The smoothing and interpolation algorithm applied uses all quadruples of consecutive existing D numbers for the least squares fitting of polynomials of second degree. The D numbers calculated by the polynomials are used to replace the D numbers of the amide hydrogens that are calculated from the directly determined D numbers of the fragments. The smoothed D number of a particular amide hydrogen is the weighted average of all the polynomial D numbers of the particular amide hydrogen. For a polynomial D number to be considered, the amino acid numbers used for the polynomial fit must include the particular amino acid number. The weighting factors decrease rapidly with the distance of the amino acid considered from the amino acid of the polynomial fit closest to the amino acid considered. It should be mentioned that the smoothing algorithm contains a routine for outlier detection.

The rationale of the smoothing and interpolation procedure described in the previous paragraph is the evidence that the D numbers of the fragments of a certain type constitute a smooth function of the fragment number. Due to the smoothing procedure, the precision of the finally employed D numbers of the amide hydrogens is significantly improved. In the case of an amide hydrogen whose D number cannot be determined directly because one of needed fragments is lacking or not suitable for analysis, the D number of that fragment is replaced by the result of the smoothing process. We believe that such interpolation is justified to complement the data by individual missing values, except where the missing D number of the amide hydrogen corresponds to an amino acid with a chemical exchange constant that is strongly deviant from the numbers of its neighbors. In this case, the experimental series of exchange rates may exhibit a discontinuity that should not be neglected by smoothing.

#### 4. Evaluation of the Exchange Kinetics

##### 4.1 Kinetic model

The widely accepted kinetic model corresponds to the following scheme:

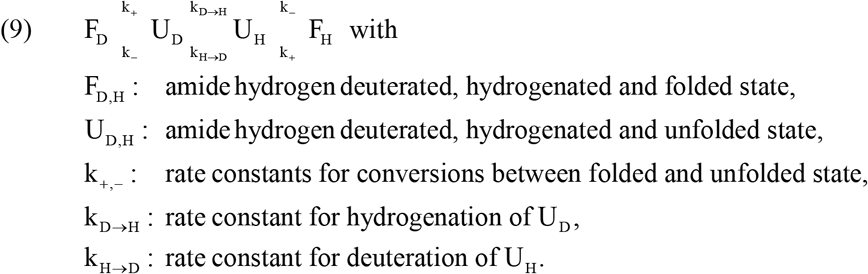

If there is no isotopic effect for the hydrogenation of U_D_ and the deuteration of U_H_, the rate constant k_H→D_ and the rate constant k_D→H_ will be proportional to the ratio of deuterated to hydrogenated exchange solvent and to the ratio of hydrogenated to deuterated exchange solvent, respectively. Hence, with the chemical exchange rate k_ch_ (exchange rate constant for fully deuterated exchange solvent) and with the above defined D_asymptotic_, the rate constants can be expressed as follows:

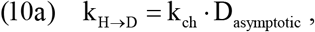

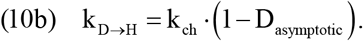

If the rate constants k_+_ and k_-_ for the local unfolding and folding of the peptide are much larger than the rate constant k_ch_, the influence of k_H→D_ and k_D→H_ on the interconversion of F_D_ and U_D_ can be neglected. This means that the concentration ratio of F_D_ and U_D_ can be approximated by the quotient k_-_/k_+_, i.e. by the value obtained for the situation where the middle part of scheme (9) is absent. The same approximation applies for the interconversion of the hydrogenated species F_H_ and U_H_.

To facilitate the solution of the dynamic system underlying the reaction scheme (9), some additional quantities are defined:

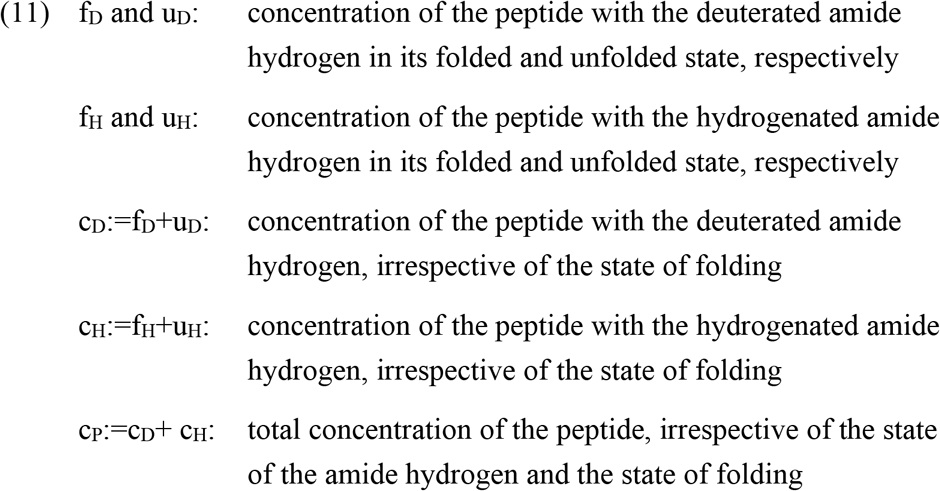

With these quantities and the assumptions outlined above, the quantities U_D_ and U_H_ needed for the formulation of the differential equation resulting from the reaction scheme (9) can be expressed by the constant overall peptide concentration c_P_ and the time dependent concentration c_D_ of the peptide with the deuterated amide hydrogen:

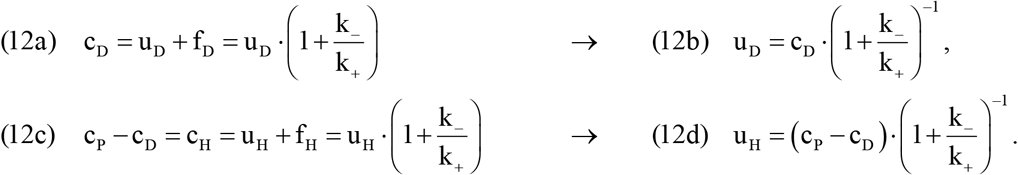

Together with the equations (10a) and (10b) and the equations (12b) and (12d), the reaction scheme (9) leads to a first order differential equation for c_D_:

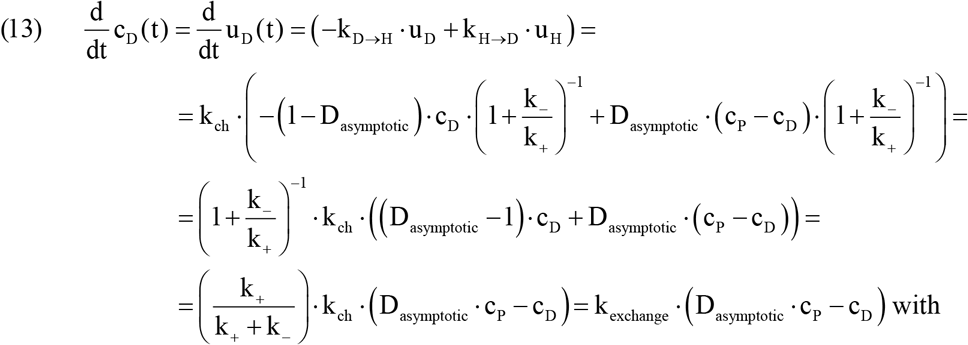

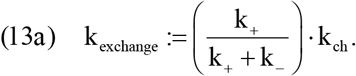

The general solution of equation (13) is the function c_D_(t) that contains a constant B needed for adjusting the initial conditions (D/H exchange: c_D_ (0)=c_p_, H/D exchange: c_D_ (0)=0):

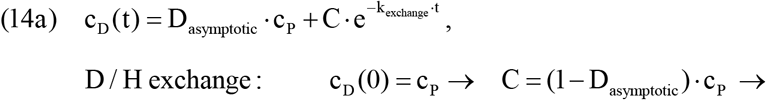

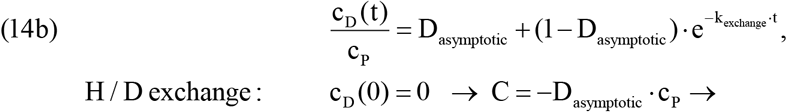

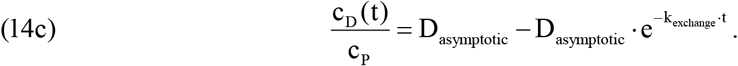

The quantity k_exchange_ equals the rate constant that is experimentally determined by fitting the time dependent D numbers of the amide hydrogens to the functions (14b) and (14c).

##### 4.2 Analysis of the exchange kinetics

The final step of the overall analysis is the analysis of the exchange kinetics. It is assumed that exchange data are available for N exchange periods t_n_ (n=1, 2,…, N). To analyze the exchange kinetics of the amide hydrogen of amino acid m, the first step consists in the determination of the mean numbers D_mean_(m,n) of the D numbers D_m_ obtained as described in section 3.3 for all the fragment types analyzed for time point n. In order to make the final results less dependent on outliers, the calculation of the mean D numbers may include outlier detection. The sensitivity of the outlier detection is decided by the user. The routine employed for outlier detection is based on the analysis of the variance of the numbers, for which the mean is calculated.

The model employed for the analysis of the kinetics is outlined in the preceding section and predicts a first order reaction that leads from the starting value D_start_ of the D number to the asymptotic D number D_asymptotic_. The experimental rate constant k_exp,m_ for the exchange of amide hydrogen m is determined by a non-linear least squares fitting routine that minimizes the quantity χ^2^ of equation (15):

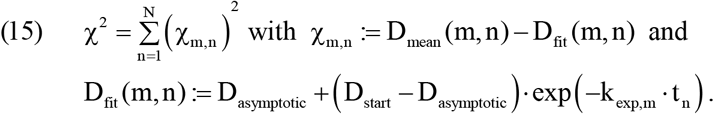

For H/D and D/H exchange, the number D_start_ amounts to 0 and 1, respectively. In case some of the numbers D_mean_(m,n) are not available due to missing fragments for certain time points, the fitting procedure is restricted to the time points with available D numbers. For the fitting to be performed, the number of time points with D numbers must not be smaller than three.

To estimate the standard error of the decimal logarithm of the experimental rate constant k_exp,m_, the standard deviation σ_m_ is calculated for the residuals χ_m,n_ obtained for the best fit value of the rate constant. The standard deviation σ_m_ is then used to perform the fitting defined in (15) for additional 2N sets of D numbers:

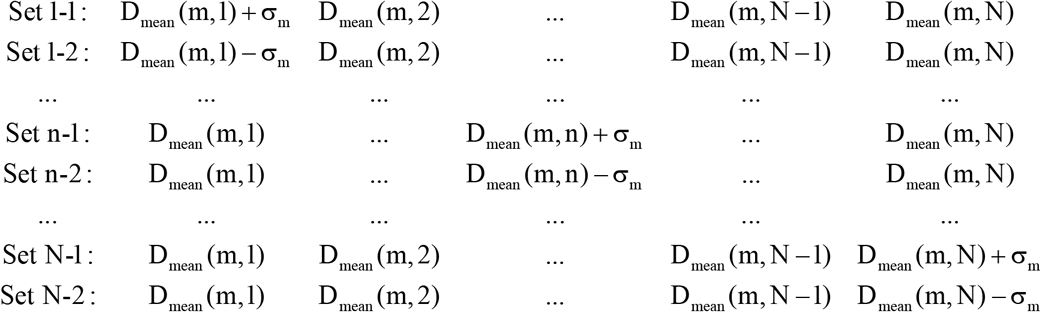

The uncertainty Δ_m,n_ of the decimal logarithm of the rate constant k_exp,m_ that is caused by the uncertainty σ_m_ of the numbers D_mean_(m,n) is estimated as the maximum of the two numbers |log[k_exp,m_]-log[k(Set n-1)]| and |log[k_exp,m_]-log[k(Set n-2)]|. The quantities k(Set n-1) and k(Set n-2) stand for the best fit rate constants obtained with the sets of D-numbers Set n-1 and Set n-2, respectively. By linear error propagation, the standard error Δlog(k_exp,m_) of the decimal logarithm of the experimental rate constant k_exp,m_ is calculated in the following way:

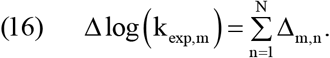

In cases where k_ch,m_ is available, the experimental rate constant k_exp,m_ can be used to calculate the difference ΔG_m_ of the Gibbs Free Energy associated with the equilibrium between the effectively folded and the effectively unfolded situation for amide hydrogen m. If the rate constant of the unfolding reaction of the peptide, which leads to an exchange competent state of amide hydrogen m, is denoted by k_m,+_, and if the rate constant of the reverse reaction that makes the amide hydrogen m not exchange competent is denoted by k_m,-_, ΔG_m_ is given by:

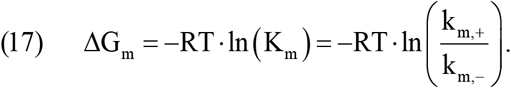

According to equation (13a), the fraction k_m,+_/(k_m,+_+ k_m,-_) can be determined from the experimental rate constant k_exp,m_ and the intrinsic rate constant k_ch,m_:

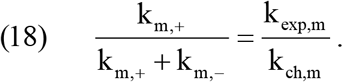

Thus, the fraction of rate constants needed for the calculation of ΔG_m_ by means of equation (17) can be calculated from the experimental rate constant k_exp,m_ and the intrinsic rate constant k_ch,m_:

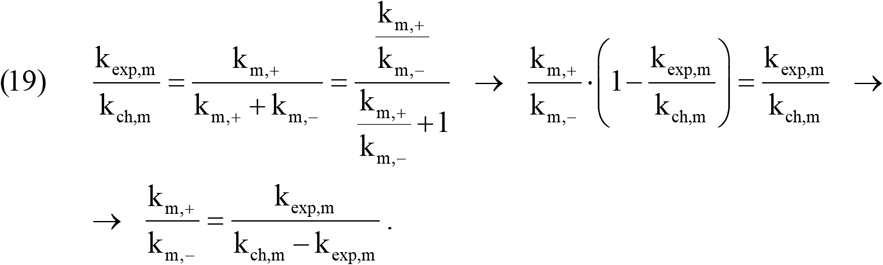

The limits of the standard confidence interval of ΔG_m_ are calculated by means of the standard error Δlog(k_exp,m_) of the decimal logarithm of k_exp,m_:

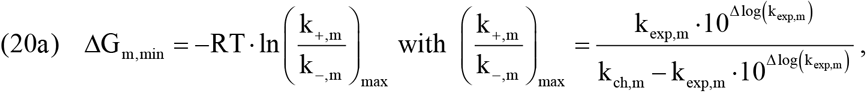

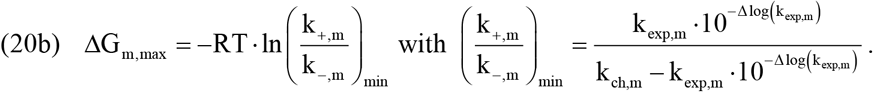

## Acknowledgements

We thank Sebastian Schuck, Harald Steiner and Dönem Avci for critical reading of the manuscript, Stefan Lichtenthaler for advice and Thomas Ruppert (ZMBH, MS facility) for MS-based cleavage site determination. This study was supported by the Deutsche Forschungsgemeinschaft (FOR2290) to DL (La699/20-1) and MKL (Le2749/1-1).

## Authors Contribution

S.S.Y. designed and performed all cell-free translocation assays, most cell-based experiments and wrote the manuscript. W.S. performed all CD and DHX experiments and analyzed data. A.L. helped determining the SPP-cleavage site. M.W. developed the *ETD Fragment Analyzer* software. D.L. and M.K.L. guided the project, designed experiments, and wrote the manuscript.

---

**Figure S1.**
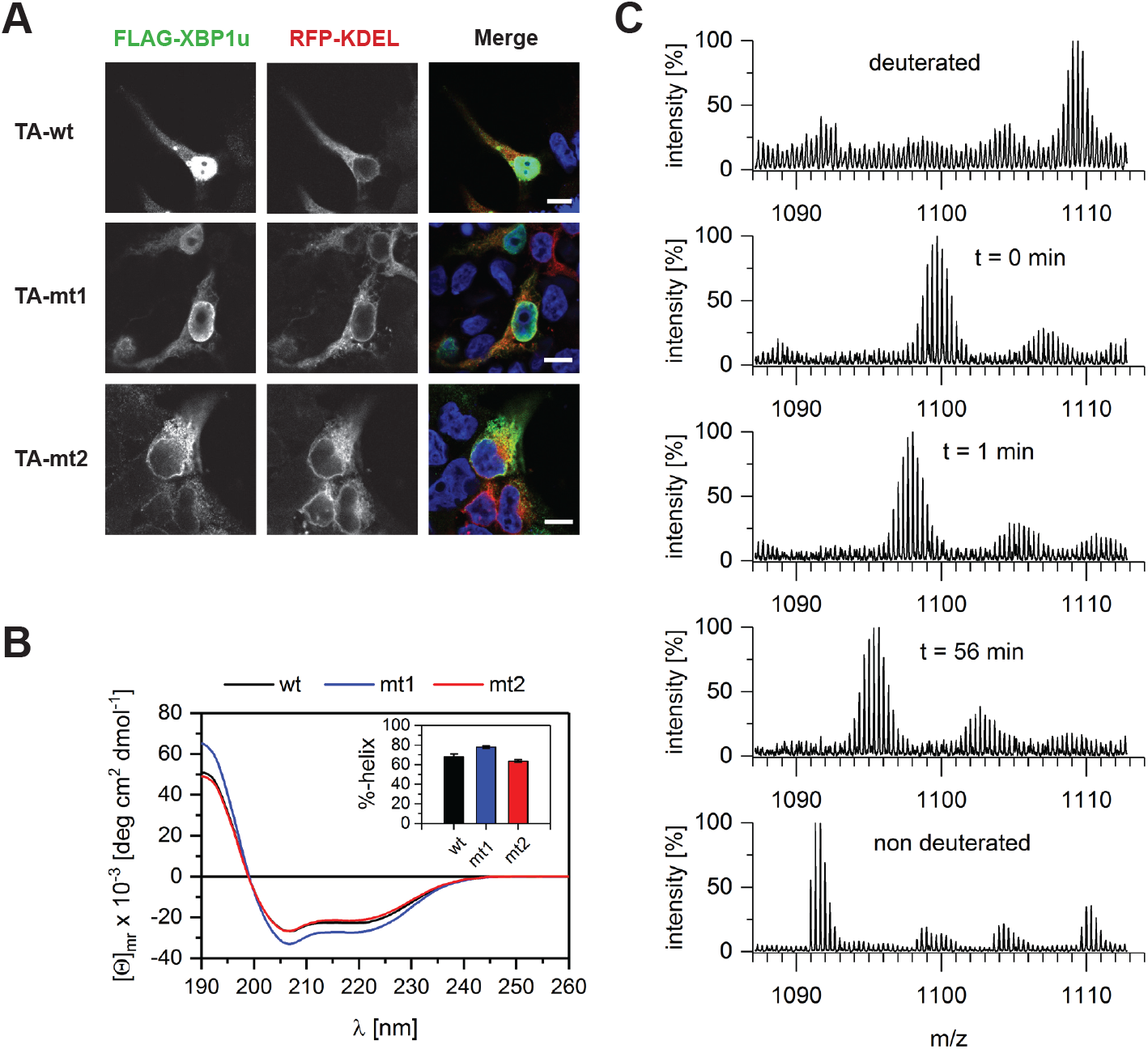
Examination of XBP1u mt1 and mt2 ER-targeting and secondary structure, related to Figure 1. **(A)** Immunofluorescent microscopy of transiently transfected Hek293T cells with FLAG-tagged tail-anchored (TA) XBP1u wt, mt1 and mt2 co-expressing the ER marker RFP-KDEL. TA-anchored XBP1u mt1 and mt2 are efficiently targeted to the ER, while TA-anchored XBP1u wt showed faint ER-staining and prominent colocalization with Hoechst-staining (shown in blue) for chromosomal DNA. Scale bar, 10 μm. **(B)** Helical content of XBP1u wt, mt1 and mt2 TM peptides determined by CD spectroscopy. CD spectra were recorded in 80% TFE/buffer, averaged, and secondary structure helix contents were calculated with CDNN (inset, n = 3, means ± SEM). **(C)** Exemplary mass spectra of the triply charged XBP1u wt ion at different time points. The spectrum at t = 0 min was recorded after exchange under stop conditions (3 min incubation on ice at pH 2.5) where only very labile deuteriums exchange. Low intensity isotopic envelopes at calculated masses ~22 Da above the dominant envelopes likely originate from Na^+^-adducts.

**Figure S2.**
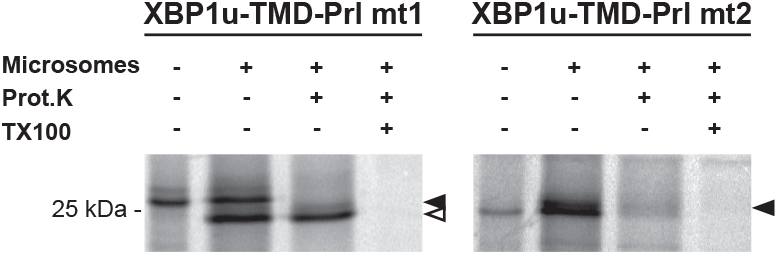
Translocation and protease protection assay of XBP1u mt1 and mt2, related to Figure 2. Translocation and protease protection assay of XBP1u-TMD-Prl mt1 and mt2 reveals that mt1 is sufficient to mediate ER insertion but mt2 is not. *In vitro* translation in the absence or presence of canine rough microsomes. Reactions were treated with Proteinase K and Triton X-100 as indicated and analyzed by SDS-PAGE and autoradiography. Filled arrowhead, pre-protein; open arrowhead, translocated Prl.

**Figure S3.**
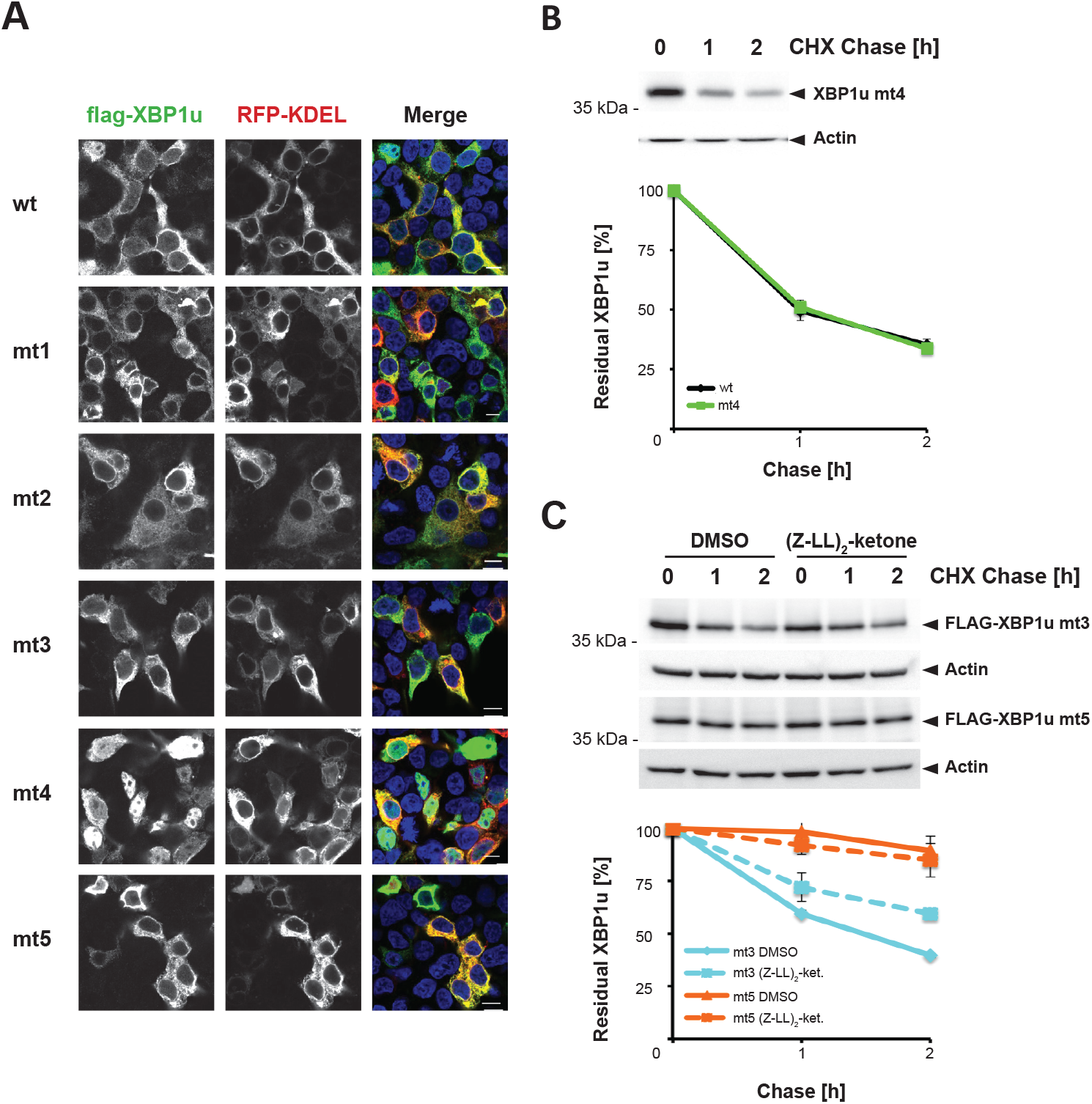
Positional effect of di-glycine motifs on ER targeting and SPP-catalyzed turnover, related to Figure 3. **(A)** XBP1u wt, mt1, mt2, mt3 and mt5 are efficiently targeted to the ER whereas mt4 predominantly localizes to the cytoplasm and nucleoplasm. Immunofluorescence microscopy of transiently transfected Hek293T cells co-expressing FLAG-tagged XBP1u full-length constructs (green) with the ER marker RFP-KDEL (red) is shown. Hoechst-staining is shown in blue. Scale bar, 10 μm. **(B)** Cycloheximide (CHX) chase analysis to determine degradation rate of XBP1u-mt4 (mean ± SEM, n=3). For comparison quantification of the degradation kinetics of XBP1u wt is shown (see Fig. 3C). **(C)** Cycloheximide (CHX) chase analysis to determine degradation rate of XBP1u mt3 and mt5 in presence of 50 μM (Z-LL)_2_-ketone (n=2) or DMSO, vehicle control.

**Figure S4.**
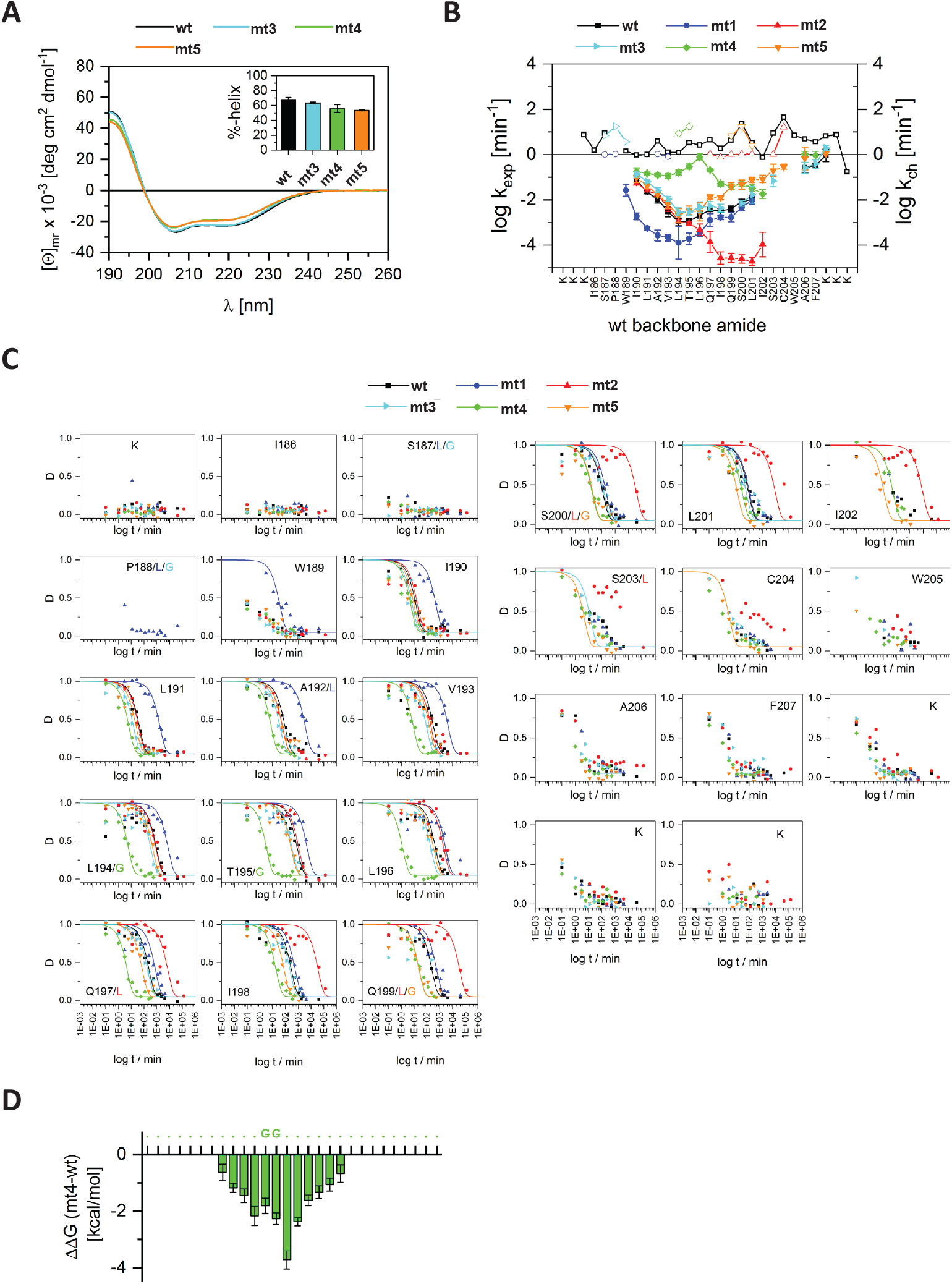
Di-glycine mutants show local TM helix destabilization, related to Figure 4. **(A)** Helical content of di-glycine mutants XBP1u mt3, mt4 and mt5 TM peptides determined by CD spectroscopy. CD spectra were recorded in 80% TFE/buffer, averaged, and secondary structure helix contents were calculated with CDNN (inset, n = 3, means ± SEM). For comparison spectrum of XBP1u wt is shown (taken from Figure S1D). **(B)** Exchange rate constants K_exp_ of individual amide deuterons. The values of K_exp_ (filled symbols, means ± SEM) were derived from exponential fits of the residue-specific DHX kinetics (given in panel C) as obtained from ETD experiments. Only those values are shown that are covered by sufficient data points. Empty symbols represent the respective chemical exchange rate constant k_ch_ that reflects the DHX kinetics in an unfolded state. **(C)** Residue-specific DHX kinetics obtained after ETD. The calculated deuterium contents D (mean values, n ≥ 3) of the respective amides are plotted against the log of the exchange period t. The identities of the wt residues (in black), along with the identities of the respective mutants (color coded) are given in the insets. Exponential fits are depicted for those kinetics that were complete enough for calculating the exchange rate constants given in panel (B) and in Table S2. **(D)** ΔΔG between wt and mt4 TM domain (compared to ΔΔG of XBP1u wt, see Fig. 4A).

**Figure S5.**
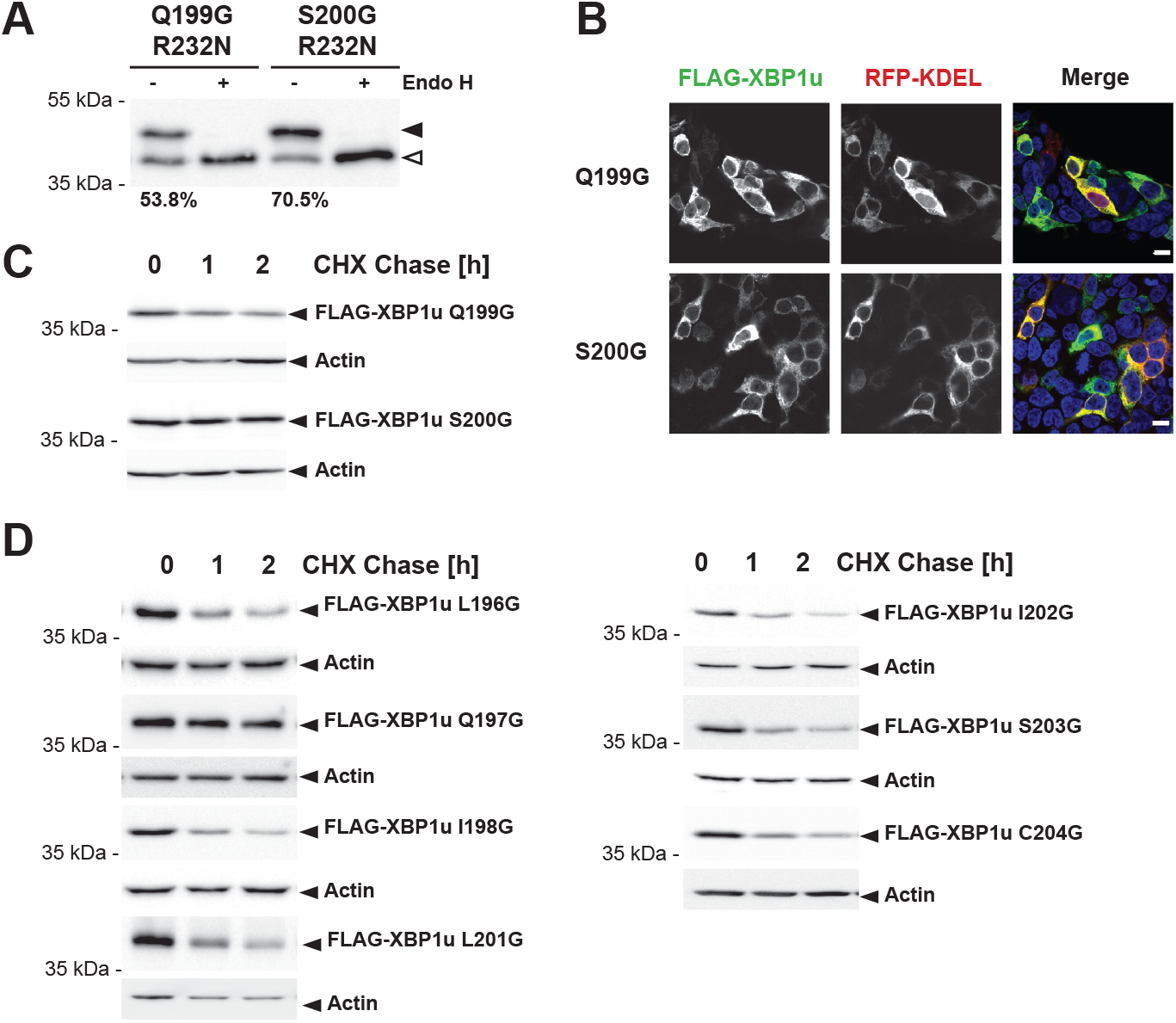
XBP1u turnover depends on residue specific features, related to Figure 5. **(A)** ER targeting and topology of XBP1u Q199G and S200G assessed by the R232N glycosylation reporter. **(B)** XBP1u Q199G and S200G localization by immunofluorescence microscopy of transiently transfected Hek293T cells co-expressing FLAG-tagged XBP1u full-length constructs (in green) with the ER marker RFP-KDEL (red) is shown. Hoechst-staining is shown in blue. Scale bar, 10 μm. **(C)** Degradation kinetics of FLAG-XBP1u Q199G and S200G in transfected Hek293T cells assessed by cycloheximide (CHX) chase and western blotting using anti-FLAG antibody (mean ± SEM, n=3). Quantification of XBP1u wt is shown for comparison (see Figure 3C). **(D)** FLAG-tagged XBP1u-wt, L196G, Q197G, I198G, L201G, I202G, S203G, C204G were transfected in Hek293T cells and their stability was assessed by cycloheximide (CHX) chase and western blotting using anti-FLAG antibody (mean ± SEM, n=3).

**Figure S6.**
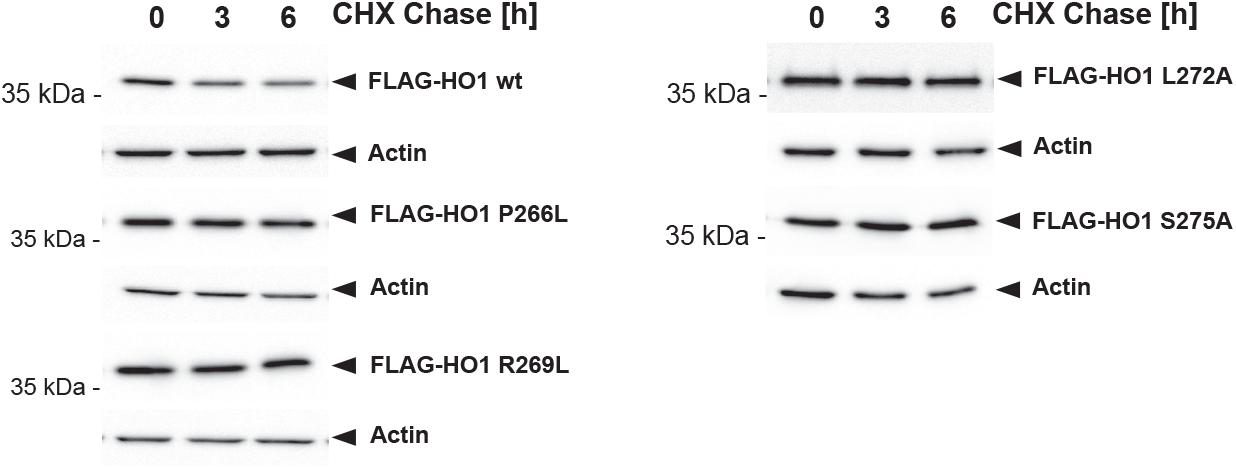
Influence of proline-266, arginine-269, leucine-272 and serine-275 on stability of HO1, related to Figure 6. FLAG-tagged HO1-wt, P266L and R269L; L272A and S275A were transfected in Hek293T cells and their stability was assessed by cycloheximide (CHX) chase and western blotting using anti-FLAG antibody (mean ± SEM, n=3).

**Table S1.** ΔG_app_ of XBP1u TM domain glycine mutants, related to Figure 3 and Figure 5.

| **XBP1u** | **Sequence** | **$\Delta G_{app}$** |
| --- | --- | --- |
| wt | ISPWILAVLTLQIQSLISCWAFWTTW | 0,07 |
| mt1 | I**LL**WIL**L**VLTLQIQSLISCWAFWTTW | -2.26 |
| mt2 | ISPWILAVLTL**L**I**LL**L**I**LCWAFWTTW | -5.765 |
| mt3 | I**GG**WILAVLTLQIQSLISCWAFWTTW | -0,27 |
| mt4 | ISPWILAV**GG**LQIQSLISCWAFWTTW | 0,766 |
| mt5 | ISPWILAVLTLQI**GG**LISCWAFWTTW | -1,222 |
| Q199G | ISPWILAVLTLQI**G**SLISCWAFWTTW | -0,986 |
| S200G | ISPWILAVLTLQIQ**G**LISCWAFWTTW | -0,158 |

**Table S2.** Numerical values of K_exp_ and ΔG, related to Figure 4.

| wt |  |  | mt1 |  |  | mt2 |  |  | mt3 |  |  | mt4 |  |  | mt5 |  |  |
| --- | --- | --- | --- | --- | --- | --- | --- | --- | --- | --- | --- | --- | --- | --- | --- | --- | --- |
| amide | log k<br>[min<sup>-1</sup>] | $\Delta G$<br>[kcal/mol] | amide | log k<br>[min<sup>-1</sup>] | $\Delta G$<br>[kcal/mol] | amide | log k<br>[min<sup>-1</sup>] | $\Delta G$<br>[kcal/mol] | amide | log k<br>[min<sup>-1</sup>] | $\Delta G$<br>[kcal/mol] | amide | log k<br>[min<sup>-1</sup>] | $\Delta G$<br>[kcal/mol] | amide | log k<br>[min<sup>-1</sup>] | $\Delta G$<br>[kcal/mol] |
| K | n.d.<sup>2</sup> |  | • | n.d. |  | • | n.d. |  | • | n.d. |  | • | n.d. |  | • | n.d. |  |
| K | n.d. |  | • | n.d. |  | • | n.d. |  | • | n.d. |  | • | n.d. |  | • | n.d. |  |
| I186 | n.d. |  | • | n.d. |  | • | n.d. |  | • | n.d. |  | • | n.d. |  | • | n.d. |  |
| S187 | n.d. |  | L | n.d. |  | • | n.d. |  | G | n.d. |  | • | n.d. |  | • | n.d. |  |
| P188 | n.d. |  | L | n.d. |  | • | n.d. |  | G | n.d. |  | • | n.d. |  | • | n.d. |  |
| W189 | n.d. |  | • | -1.58 ± 0.27 | 2.4 ± 0.4 | • | n.d. |  | • | n.d. |  | • | n.d. |  | • | n.d. |  |
| I190 | -1.15 ± 0.08<sup>3</sup> | 1.5 ± 0.1<sup>4</sup> | • | -2.73 ± 0.15 | 3.6 ± 0.2 | • | -1.28 ± 0.05 | 1.6 ± 0.1 | • | -0.87 ± 0.13 | 1.0 ± 0.2 | • | -0.75 ± 0.16 | 0.8 ± 0.3 | • | -1.01 ± 0.23 | 1.2 ± 0.4 |
| L191 | -1.65 ± 0.07 | 2.2 ± 0.1 | • | -3.26 ± 0.08 | 4.4 ± 0.1 | • | -1.58 ± 0.04 | 2.1 ± 0.1 | • | -1.19 ± 0.06 | 1.5 ± 0.1 | • | -0.83 ± 0.08 | 1.0 ± 0.1 | • | -1.39 ± 0.11 | 1.8 ± 0.2 |
| A192 | -1.98 ± 0.16 | 3.4 ± 0.2 | L | -3.57 ± 0.24 | 4.8 ± 0.3 | • | -1.78 ± 0.08 | 3.2 ± 0.1 | • | -1.58 ± 0.12 | 2.9 ± 0.2 | • | -0.91 ± 0.07 | 2.0 ± 0.1 | • | -1.76 ± 0.12 | 3.2 ± 0.2 |
| V193 | -2.52 ± 0.23 | 3.5 ± 0.3 | • | -3.68 ± 0.13 | 4.8 ± 0.2 | • | -2.33 ± 0.08 | 3.3 ± 0.2 | • | -1.97 ± 0.11 | 2.8 ± 0.2 | • | -0.95 ± 0.10 | 1.4 ± 0.1 | • | -2.20 ± 0.18 | 3.1 ± 0.2 |
| L194 | -2.98 ± 0.17 | 4.1 ± 0.2 | • | -3.89 ± 0.72 | 5.3 ± 1.0 | • | -2.94 ± 0.08 | 4.1 ± 0.1 | • | -2.53 ± 0.18 | 3.5 ± 0.2 | G | -0.79 ± 0.12 | 2.3 ± 0.2 | • | -2.68 ± 0.30 | 3.7 ± 0.4 |
| T195 | -2.93 ± 0.10 | 4.6 ± 0.1 | • | -3.72 ± 0.26 | 5.7 ± 0.3 | • | -3.05 ± 0.08 | 4.8 ± 0.1 | • | -2.53 ± 0.17 | 4.1 ± 0.2 | G | -0.53 ± 0.11 | 2.4 ± 0.2 | • | -2.46 ± 0.19 | 4.0 ± 0.2 |
| L196 | -2.69 ± 0.15 | 4.2 ± 0.2 | • | -3.47 ± 0.14 | 5.2 ± 0.2 | • | -3.33 ± 0.26 | 5.0 ± 0.3 | • | -2.32 ± 0.24 | 3.7 ± 0.3 | • | -0.11 ± 0.13 | 0.5 ± 0.3 | • | -2.53 ± 0.40 | 4.0 ± 0.5 |
| Q197 | -2.44 ± 0.08 | 4.1 ± 0.1 | • | -2.90 ± 0.28 | 4.8 ± 0.4 | L | -3.86 ± 0.46 | 5.2 ± 0.6 | • | -2.28 ± 0.20 | 3.9 ± 0.3 | • | -0.69 ± 0.07 | 1.8 ± 0.1 | • | -1.87 ± 0.09 | 3.4 ± 0.1 |
| I198 | -2.48 ± 0.09 | 3.7 ± 0.1 | • | -2.77 ± 0.10 | 4.1 ± 0.1 | • | -4.57 ± 0.28 | 6.0 ± 0.4 | • | -2.36 ± 0.17 | 3.5 ± 0.2 | • | -1.28 ± 0.11 | 2.1 ± 0.1 | • | -1.91 ± 0.09 | 2.9 ± 0.1 |
| Q199 | -2.40 ± 0.12 | 4.1 ± 0.2 | • | -2.77 ± 0.20 | 4.6 ± 0.3 | L | -4.57 ± 0.20 | 6.1 ± 0.3 | • | n.d. |  | • | -1.41 ± 0.11 | 2.7 ± 0.1 | G | -1.39 ± 0.16 | 3.0 ± 0.2 |
| S200 | -2.08 ± 0.11 | 4.6 ± 0.1 | • | -2.33 ± 0.16 | 5.0 ± 0.2 | L | -4.61 ± 0.19 | 6.2 ± 0.3 | • | -2.15 ± 0.13 | 4.7 ± 0.2 | • | -1.28 ± 0.13 | 3.6 ± 0.2 | G | -1.32 ± 0.20 | 3.4 ± 0.3 |
| L201 | -1.98 ± 0.18 | 3.4 ± 0.2 | • | -2.03 ± 0.18 | 3.4 ± 0.2 | • | -4.72 ± 0.18 | 6.4 ± 0.2 | • | -1.79 ± 0.37 | 3.1 ± 0.5 | • | -1.48 ± 0.14 | 2.7 ± 0.2 | • | -1.12 ± 0.12 | 2.0 ± 0.2 |
| I202 | n.d. |  | • | n.d. |  | • | -3.98 ± 0.53 | 5.2 ± 0.7 | • | n.d. |  | • | -1.74 ± 0.20 | 2.2 ± 0.3 | • | -1.07 ± 0.16 | 1.2 ± 0.2 |
| S203 | n.d. |  | • | n.d. |  | L | n.d. |  | • | -1.16 ± 0.26 | 2.8 ± 0.4 | • | n.d. |  | • | -0.70 ± 0.27 | 2.2 ± 0.4 |
| C204 | n.d. |  | • | n.d. |  | • | n.d. |  | • | n.d. |  | • | n.d. |  | • | -0.54 ± 0.11 | 2.9 ± 0.2 |
| W205 | n.d. |  | • | n.d. |  | • | n.d. |  | • | n.d. |  | • | n.d. |  | • | n.d. |  |
| A206 | -0.58 ± 0.24 | 1.7 ± 0.3 | • | n.d. |  | • | n.d. |  | • | -0.59 ± 0.25 | 1.7 ± 0.4 | • | 0.01 ± 0.14 | 0.8 ± 0.2 | • | -0.17 ± 0.51 | 1.1 ± 0.9 |
| F207 | -0.49 ± 0.10 | 1.4 ± 0.1 | • | n.d. |  | • | n.d. |  | • | -0.41 ± 0.19 | 1.2 ± 0.3 | • | -0.04 ± 0.18 | 0.6 ± 0.4 | • | n.d. |  |
| K | -0.04 ± 0.28 | 1.1 ± 0.5 | • | n.d. |  | • | n.d. |  | • | -0.29 ± 0.11 | 0.5 ± 0.2 | • | n.d. |  | • | 0.03 ± 0.10 | 0.9 ± 0.2 |
| K | n.d. |  | • | n.d. |  | • | n.d. |  | • | n.d. |  | • | n.d. |  | • | n.d. |  |
| K | n.d. |  | • | n.d. |  | • | n.d. |  | • | n.d. |  | • | n.d. |  | • | n.d. |  |

<sup>1</sup> Identity of the TM domain peptide
<sup>2</sup> n.d. = not determined
<sup>3</sup> $k_{\text{exp}}$ from exponential fit of means ± SE
<sup>4</sup> $\Delta G$ with confidence interval

**Table S3. Primers used in this study.**

See separate file.

